# Zoonotic avian influenza viruses evade human BTN3A3 restriction

**DOI:** 10.1101/2022.06.14.496196

**Authors:** Rute Maria Pinto, Siddharth Bakshi, Spyros Lytras, Mohammad Khalid Zakaria, Simon Swingler, Julie C Worrell, Vanessa Herder, Margus Varjak, Natalia Cameron-Ruiz, Mila Collados Rodriguez, Mariana Varela, Arthur Wickenhagen, Colin Loney, Yanlong Pei, Joseph Hughes, Elise Valette, Matthew L Turnbull, Wilhelm Furnon, Kerrie E Hargrave, Quan Gu, Lauren Orr, Aislynn Taggart, Chris Boutell, Finn Grey, Edward Hutchinson, Paul Digard, Isabella Monne, Sarah K Wootton, Megan K L MacLeod, Sam J Wilson, Massimo Palmarini

## Abstract

Cross-species transmission of avian influenza A viruses (IAVs) into humans could represent the first step of a future pandemic^1^. Multiple factors limiting the spillover and adaptation of avian IAVs in humans have been identified, but they are not sufficient to explain which virus lineages are more likely to cross the species barrier^1,2^. Here, we identified human BTN3A3^3^ (butyrophilin subfamily 3 member A3) as a potent inhibitor of avian but not human IAVs. We determined that BTN3A3 is constitutively expressed in human airways and its antiviral activity evolved in primates. We show that BTN3A3 restriction acts at the early stages of virus replication by inhibiting avian IAV vRNA transcription. We identified residue 313 in the viral nucleoprotein (NP) as the genetic determinant of BTN3A3 sensitivity (313F, or rarely 313L in avian viruses) or evasion (313Y or 313V in human viruses). However, several serotypes of avian IAVs that spilled over into humans in recent decades evade BTN3A3 restriction. In these cases, BTN3A3 evasion is due to substitutions (N, H or Q) in NP residue 52 that is adjacent to residue 313 in the NP structure^4^. Importantly, we identified more than 150 avian IAV lineages with a BTN3A3-resistant genotype. In conclusion, sensitivity or resistance to BTN3A3 is another factor to consider in the risk assessment of the zoonotic potential of avian influenza viruses.

## Main Text

Influenza A viruses (IAVs) cause a substantial global health burden and circulate both in humans and animal species including domestic poultry, pigs, dogs and horses. Wild aquatic birds are the main natural reservoir of IAV^5^. Ducks, shorebirds, gulls and other waterbirds harbour 16 haemagglutinin (HA) and 9 neuraminidase (NA) subtypes^6^. IAV from wild bird reservoirs can infect terrestrial birds such as chickens, turkeys, quail and other gallinaceous species, or domestic waterfowl^5^. The population density of domestic birds allows opportunities for IAVs to spill over into susceptible mammals, including humans. Avian IAVs can also further reassort (i.e. exchange genome segments) with viruses established in susceptible species resulting in a rich genetic diversity for this group of viruses. In humans, the influenza pandemics of 1918, 1957, 1968 and 2009 were all caused by viruses containing at least some genomic segments of avian origin^7,8^.

Occasionally, transmission of avian IAVs into humans occurs and may result in severe or even lethal disease^9^. For example, in 2013 an H7N9 variant, resulting from multiple reassortment events of different avian viruses^10^, caused more than 600 deaths in humans^11^. These spillover events are not typically followed by extensive human-to-human transmission chains, but they are a risk to global health as they could enable the first step towards adaptation to the human host and the generation of pandemic IAV strains^1^.

Multiple barriers have been identified that hamper avian IAVs transmission and adaptation in humans^12^. These include virus haemagglutinin (HA) receptor binding specificity^13^, the pH that activates HA^14,15^, the efficiency of the virus polymerase^16^, the length of the viral neuraminidase (NA) stalk^17,18^, and sensitivity to the host antiviral factor Mx1/MxA^19,20^. For example, the unusually high number of human infections caused by H7N9, has been correlated with changes in the viral glycoproteins, and the polymerase complex^21^. Correlation between disease severity induced by H7N9 and certain human Mx1 alleles has also been described in GWAS studies^22^.

Despite the progress made in the last two decades, there is an incomplete understanding of what allows certain avian IAV subtypes/lineages to spill over in humans. In this study, we aimed to identify (i) host genetic barriers to avian IAVs replication in human cells and (ii) IAV genetic signatures that can be directly correlated with their zoonotic potential.

### Human BTN3A3 potently and specifically restrict avian IAVs

We focused on the host type-I interferon (IFN) response as it is one of the key host antiviral innate immune mechanisms and a barrier for virus cross-species transmission^23^. IFN acts through the activation of hundreds of interferon stimulated genes (ISGs), some of which have antiviral properties^23^. Hence, we first aimed to identify human ISGs that contribute to IAVs host tropism. We performed arrayed expression screening^24,25^ of 752 human and macaque ISGs (Fig. 1a) using three different recombinant IAV strains, each tagged with GFP: A/Puerto Rico/8/1934 (PR8, a human laboratory adapted H1N1 strain), A/California/04/2009 (Cal04, a mouse-adapted pdm09 H1N1 strain) and A/mallard/Netherlands/10-Cam/1999 (Mallard, a H1N1 avian strain). These screens (Fig. 1b, Extended Data Fig. 1) identified ISGs previously shown to be antiviral against human-origin IAVs, such as IFITM2, IFITM3 and Mx1^26–29^. However, two other ISGs, BTN3A1 and BTN3A3, inhibited the avian virus (Mallard), but not the mammalian viruses (PR8 or Cal04).

**Figure 1:**
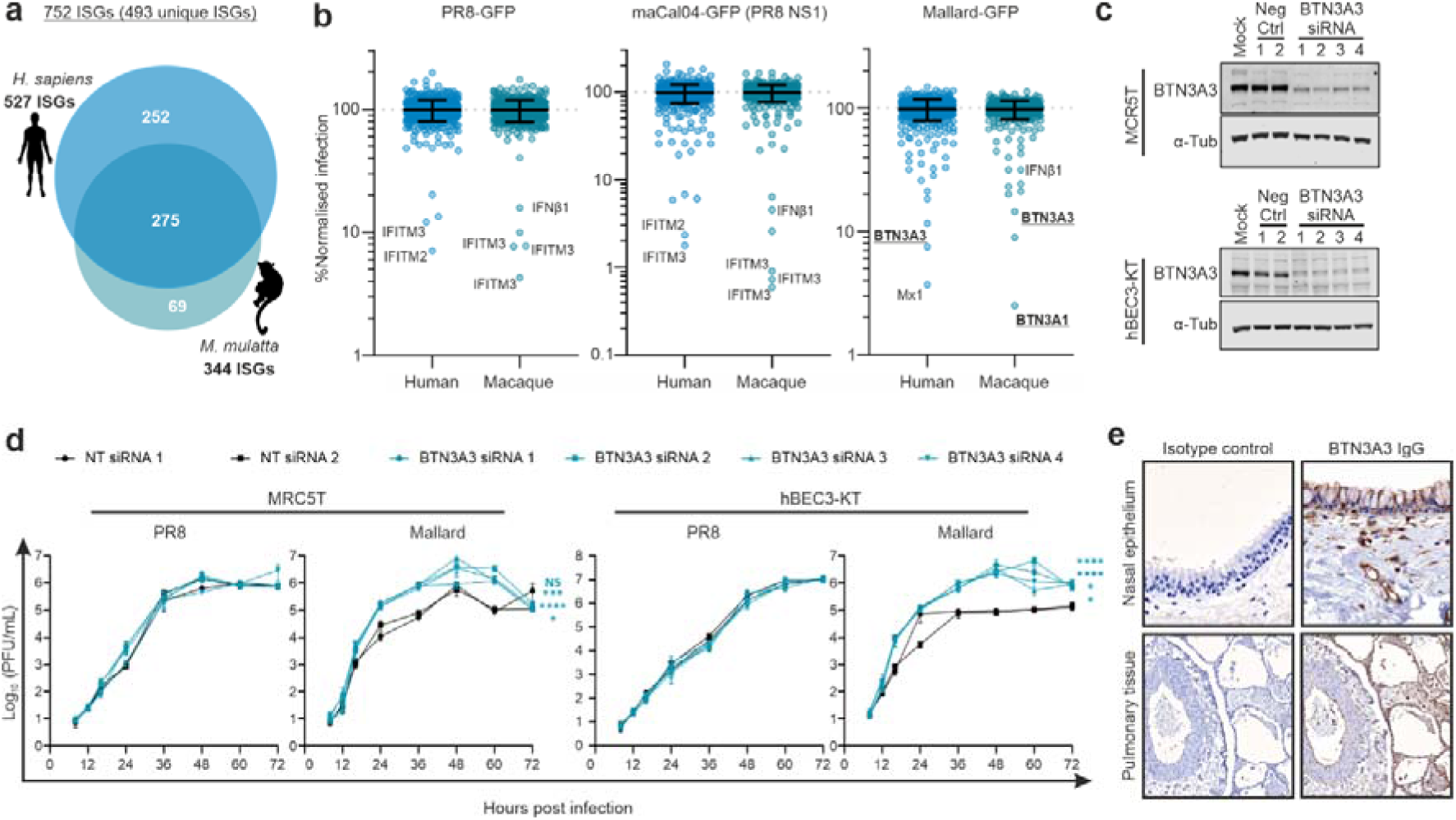
Arrayed ISG libraries screens identify BTN proteins as restriction factors for avian IAV. **a** Schematic representation of the ISG libraries used in the screens. **b**, Graphs showing the results of normalised infections (% median of overall library) of cells expressing individual ISGs with the indicated viruses. ISGs restricting virus replication are labelled. **c**, Western blotting showing expression and effective siRNA-driven knock-down of BTN3A3 in MRC5T and hBEC3-KT. Cells were transfected with scrambled (Neg ctrl) or BTN3A3-targetting siRNAs, and protein levels in the resulting cell lysates were assessed by western blotting. α-Tubulin was used as loading control. **d**, Replication kinetics of PR8 and Mallard in siRNA-treated MRC5T and hBEC3-KT. Cells were infected with a MOI of 0.001 and supernatants were collected at the mentioned times post-infection. Infectious viral titres were determined by plaque assay. Data are mean +/- standard error of the mean (SEM) of 3 independent experiments. Statistical significance between groups was measured by a 2-way ANOVA. Comparisons were made between area under the curve of the different BTN3A3 siRNA treatment conditions and the average of the two Neg ctrl. NS-Non-significant, *p≤0.05, **p≤0.01, ***p≤0.001, ****p≤0.0001. **e**, Micrographs of human nasal epithelium and lung sections subjected to immunohistochemistry. Cells expressing BTN3A3 (shown in brown) are found along the entire nasal epithelium and in endothelial cells, immune cells as well as pneumocytes.

In order to validate these data, we carried out loss of function experiments. We detected constitutive expression of BTN3A3 in primary immortalised human foetal lung fibroblasts (MRC5T) and human bronchial epithelial cells (hBEC3-KT) (Fig. 1c). siRNA-mediated knockdown of BTN3A3 in these cells resulted in improved replication of the avian Mallard strain, while it did not affect the growth kinetics of PR8 (Fig. 1d). Equivalent knockdown experiments targeting BTN3A1 did not show a significant difference in avian IAVs replication (Extended Data Fig. 2).

By immunohistochemistry we showed that BTN3A3 is constitutively expressed in the respiratory tract of healthy/uninfected individuals, both in upper and lower respiratory tract. We saw BTN3A3 expression in ciliated cells within the nasal epithelium and bronchioli, type I and II alveolar epithelial cells and alveolar macrophages (Fig. 1e). Indeed, available data from human transcriptomic profiles (GTEx consortium) suggest that the lungs have the second highest expression levels of BTN3A3 across 30 tissues analysed (Extended Data Fig. 3).

The butyrophilin gene superfamily has undergone complex duplication events over its evolutionary history with subfamily 3 comprising of three primate-specific paralogues: BTN3A1-3^3,30^. We tested the antiviral activity of overexpressed human BTN3A1, BTN3A2 and BTN3A3 in A549 cells (Fig. 2a). Infection of BTN3-overexpressing cells using GFP-tagged and untagged viruses showed that PR8 was not sensitive to any of the human BTN3 proteins. However, BTN3A1 and BTN3A3 were successful at restricting Mallard (the latter being more effective at restriction) whilst BTN3A2 showed no antiviral effects (Fig. 2b-c). Next, we assessed the antiviral effects of the BTN3 proteins against a wider panel of IAV strains. These included human laboratory-adapted viruses, human clinical isolates and various avian influenza viruses isolated from divergent bird species. None of the human viruses tested were sensitive to any of the BTN3A proteins (Fig. 2d, left panels). However, the replication of all avian viruses was restricted approximately 10-fold by BTN3A1. Restriction of avian viruses by BTN3A3 was even greater with viral titres barely reaching the limit of detection and showing a decrease of up to 100,000-fold (Fig. 2d, right panels).

**Figure 2:**
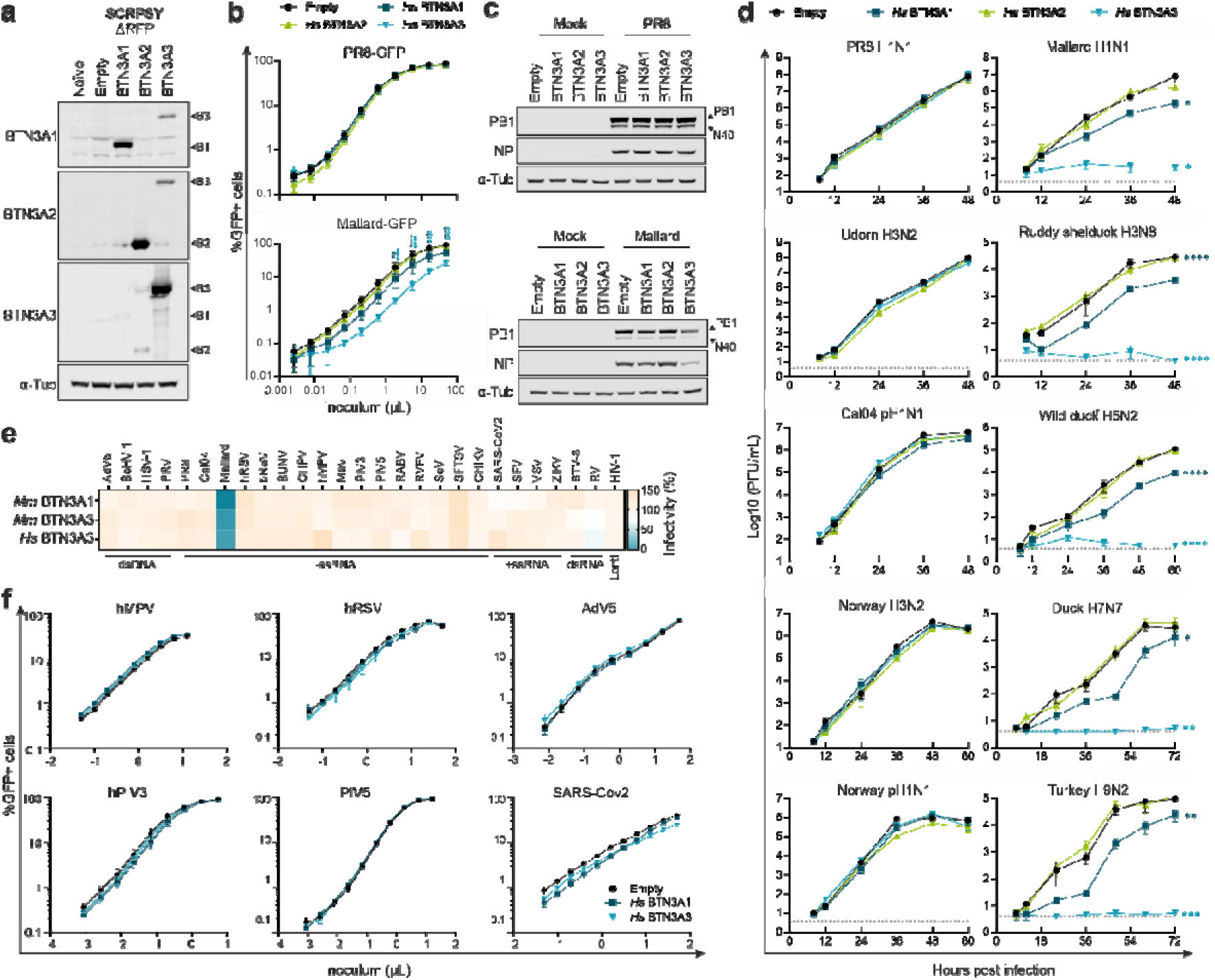
BTN3A1 and BTN3A3 restrict replication of avian but not human IAV. **a,** Western blotting of cell lysates obtained from A549 cells stably expressing human BTN3A1, BTN3A2 or BTN3A3. Due to their sequence and domain similarity, cross-detection of BTN3A3 was observed with the BTN3A1 and BTN3A2 antibodies, while BTN3A2 is detected also using anti-BTN3A3 antibodies. **b**, BTN-overexpressing cells were challenged with serial dilutions of GFP-tagged PR8 or Mallard for 10h. Upon trypsinisation and fixation, percentage of GFP-positive cells was measured by flow cytometry. Data are mean +/- SEM of 3 independent experiments. Statistical significance between groups was measured by a 1-way ANOVA for each virus dose. Comparisons were made between each BTN-expressing cells and empty control. *p≤0.05, **p≤0.01, ***p≤0.001, ****p≤0.0001. **c**, A549 cells as in (a) and (b) were infected with PR8 or Mallard WT viruses at a MOI 3 for 6h. Viral protein expression was measured by detection of PB1 (and PB1-N40) and NP by western blot. **d**, Replication kinetics of a panel of human (left panels) and avian (right panels) viruses in A549-Empty and BTN3A1 or BTN3A3-expressing cells. Cells were infected at an MOI of 0.001. Supernatants were collected at the indicated timepoints post-infection and infectious virus titres were obtained by plaque assay. A/Puerto Rico/8/1934 H1N1 (PR8 H1N1), A/Udorn/307/1972 H3N2 (Udorn H3N2), A/California/04/2009 H1N1 (Cal04 H1N1), A/Norway/2433 2018 H3N2 (Norway H3N2), A/Norway/3275/ 2018 H1N1 (Norway H1N1), A/mallard/Netherlands/10-Cam/1999 H1N1 (Mallard H1N1), A/ruddy shelduck/Mongolia/963V/2009 H3N8 (Ruddy shelduck H3N8), A/wild-duck/Italy/17VIR6926-1/2017 H5N2 (Wild duck H5N2), A/duck/Italy/18VIR4932-2/2018 H7N7 (Duck H7N7), A/turkey/Italy/16VIR8643-54/2016 H9N2 (Turkey H9N2). Data represent mean +/- SEM of 3 independent experiments. Statistical significance between groups was measured by a 2-way ANOVA. Comparisons were made between area under the curve of the different BTN-expressing cells and empty control. *p≤0.05, **p≤0.01, ***p≤0.001, ****p≤0.0001. **e**, Heatmap showing ISG screens (similar to Fig. 1b) and normalised infection of cells expressing BTN3A1 and BTN3A3 for a panel of viruses. Details can be found in Materials and Methods. **f**, Human BTN3A1 and BTN3A3 overexpressing A549s were challenged with different volumes of the GFP-tagged viruses indicated in the figure. Cells were fixed at 10 hours for PR8 and Mallard and 16 hours post infection for the remaining viruses. Percentage of GFP-positive cells was measured by flow cytometry. Data are mean +/- SEM of 3 independent experiments. Statistical significance between groups was measured by a 1-way ANOVA for each virus dose. Comparisons were made between each BTN-expressing cells and empty control and no significant differences were found.

We further tested the antiviral specificity of BTN3A1 and BTN3A3 against an additional 24 viruses from a selection of genera, including both dsDNA, ssRNA and dsRNA viruses, including both enveloped and non-enveloped viruses. Of all the screened viruses, Mallard was the only virus substantially inhibited by BTN3A1 and BTN3A3 (Fig. 2e). Further validation assays against a panel of human respiratory viruses confirmed the specificity of this restriction factor against avian IAVs (Fig. 2f).

### BTN3A3 antiviral activity likely evolved after the split between old and new world monkeys

We then examined the origin of anti-avian IAV activity in the BTN3 gene family. Phylogenetic analysis of the BTN3A genes of the *Haplorrhini* sub-order (tarsier, monkey, apes, and humans) indicated that BTN3A1-3 originated through two successive duplications after the split between the new world monkeys lineage (*Platyrrhini*) and the old world monkeys and apes lineage (*Catarrhini*) around 40 – 44 million years ago (Fig. 3a)^31^. Domain detection analysis showed that the majority of BTN3A1 and BTN3A3 genes have a consistent domain organisation with one set of N-terminal IgV and IgC domains followed by a PRYSPRY domain, while BTN3A2 genes have lost their PRYSPRY domain (with the exception of *Nomascus leucogenys* BTN3A2).

**Figure 3:**
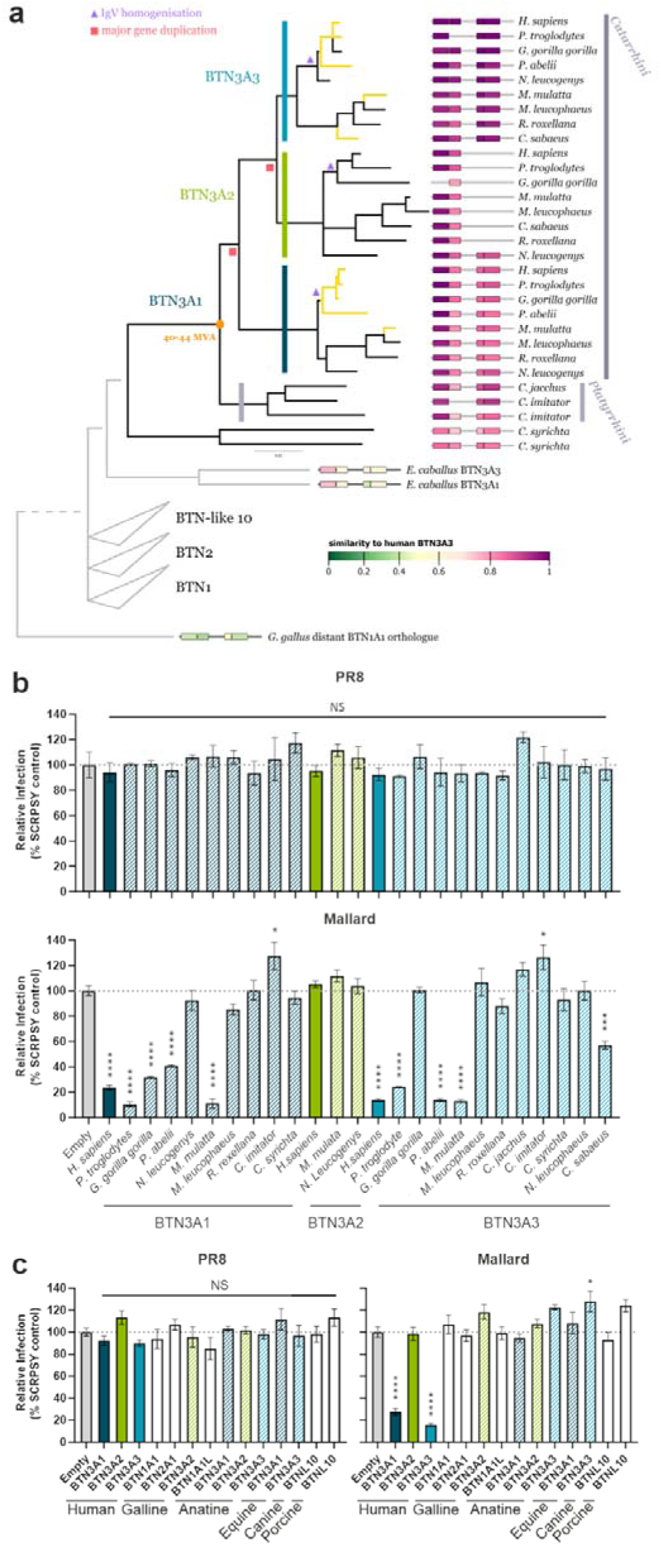
Evolution of antiviral activity of BTNs. **a**, Maximum likelihood phylogeny of concatenated protein domain coding sequences the Haplorrhini BTN3 genes (K2P+G4 substitution model). Nodes with bootstrap support below 60 have been collapsed. Branches confirmed to have anti-avian IAV activity (see below) are highlighted in yellow. Relation to more distant tested homologues and orthologous/paralogous gene families are shown as a schematic in grey. IgV homogenisation events, major gene duplications and gene subfamilies are annotated on the phylogeny. Presence of each of the four protein domains (IgV, IgC, PRY and SPRY) is annotated on the right of each tree tip and coloured by pairwise amino acid similarity to the respective domain of the human BTN3A3. Species names and taxonomic classification is annotated on the right. The median divergence time between *Catarrhini* and *Platyrrhini* was retrieved from timetree. **b-c**, A549 cells w re transiently transduced with SCRPSY lentiviruses expressing the indicated BTN proteins and challenged with PR8- or Mallard-GFP. 8 h post infection, the percentage of GFP-positive cells was measured by flow cytometry. Data are mean +/- SEM of 2 independent experiments. Statistical significance between groups was measured by a 1-way ANOVA. Comparisons were made between each BTN-expressing cells and empty control. NS-Non-significant, *p≤0.05, **p≤0.01, ***p≤0.001, ****p≤0.0001.

Transient transduction of A549 cells with BTN3A-expressing lentiviruses followed by viral challenge showed, that PR8 was not restricted by any *Haplorrhini* BTN3 proteins (Fig. 3b). Mallard was inhibited specifically by the *Catarrhini* (old world monkeys and apes) BTN3A1/3 genes but not by *Platyrrhini* (new world monkeys) or *Tarsiiformes* (tarsier) BTN3 genes (Fig. 3b). This suggests that the antiviral phenotype of BTN3A3 was gained after the *Platyrrhini*-*Catarrhini* split, consistent with the two duplication events. Humans, chimpanzees (*Pan troglodytes*), gorillas (*Gorilla gorilla*), orangutans (*Pongo abelii*), macaques (*Macaca mulatta*) and green monkeys (*Chlorocebus sabaeus*) all have at least one BTN3A1/3 gene capable of inhibiting Mallard viral replication (Fig. 3b). We also found that the closest orthologues to BTN3 from mammalian species susceptible to IAVs (canine – *Canis lupus familiaris*, equine - *Equus caballus*, and porcine - *Sus scrofa*), distant galline (*Gallus gallus*) and anatine (*Anas platyrhynchos*) BTN1 orthologues, and the human paralogues of the butyrophilin superfamily did not inhibit PR8 or Mallard, supporting a *Catarrhini* origin of BTN3 antiviral activity (Fig. 3b, Extended Data Fig. 4). Our phylogenetic analysis which considers each *Haplorrhini* BTN3 protein domain individually is consistent with previously documented recombination and homogenisation of the IgV domain within the BTN3 *Catarrhini* gene subfamily^32^ (Extended Data Fig. 5). This incongruence in the evolutionary history of these genes could explain the multiple gains/losses of antiviral function.

### NP residue 313 is a key determinant of sensitivity/resistance to BTN3A3

To identify the viral genetic determinants of BTN3A3 sensitivity or evasion, we engineered reassortants between PR8 and Mallard and tested their ability to replicate in BTN3A3-overexpressing MDCK cells (Extended data Fig. 6a). All PR8-based reassortants formed plaques as efficiently as wild type virus in BTN3A3-expressing cells apart from those containing Mallard segment 5 (Extended data Fig. 6b). The opposite phenotype was seen in Mallard-based reassortants in which only viruses containing PR8-segment 5 (specifically Mallard with PR8 containing segments encoding for the viral ribonucleoproteins) managed to successfully form plaques in BTN3A3-overexpressing MDCK. We were unable to rescue the single segment reassortant Mallard 7:1 PR8 Segment 5. Hence, we engineered the reciprocal segment 5 reassortants between Mallard and Cal04. We assessed virus propagation in A549-Empty and BTN3A3 expressing cells (A549-BTN3A3), and confirmed that Mallard segment 5 conferred a BTN3A3 sensitive phenotype (Fig. 4a). Segment 5 is monocistronic and encodes for the viral nucleoprotein (NP)^33^. To identify the amino acid residue/s in NP determining the sensitivity or resistance to BTN3A3, we compared the NP sequences of the five human and five avian IAVs tested in Fig. 3d. Human and avian NP sequences have conserved differences in residues 33, 100, 136, 313, 351, 353 and 357. Of these, positions 33, 100, 313 and 357 have been previously associated with avian-to-human transmission^34^. We rescued PR8 or Mallard single NP mutants and assessed their replication in A549-Empty or A549-BTN3A3. Using a PR8 background, we observed no reduction in virus yields in A549-BTN3A3 for any mutants except for PR8-Y313F. Conversely, on a Mallard-background the R100V mutant resulted in partial loss of BTN3A3 sensitivity but F313Y evaded BTN3A3 restriction almost completely (Fig. 4b; Extended Data Fig. 7). Therefore, we conclude that amino acid residue 313 is a key determinant of BTN3A3 sensitivity/evasion.

**Figure 4:**
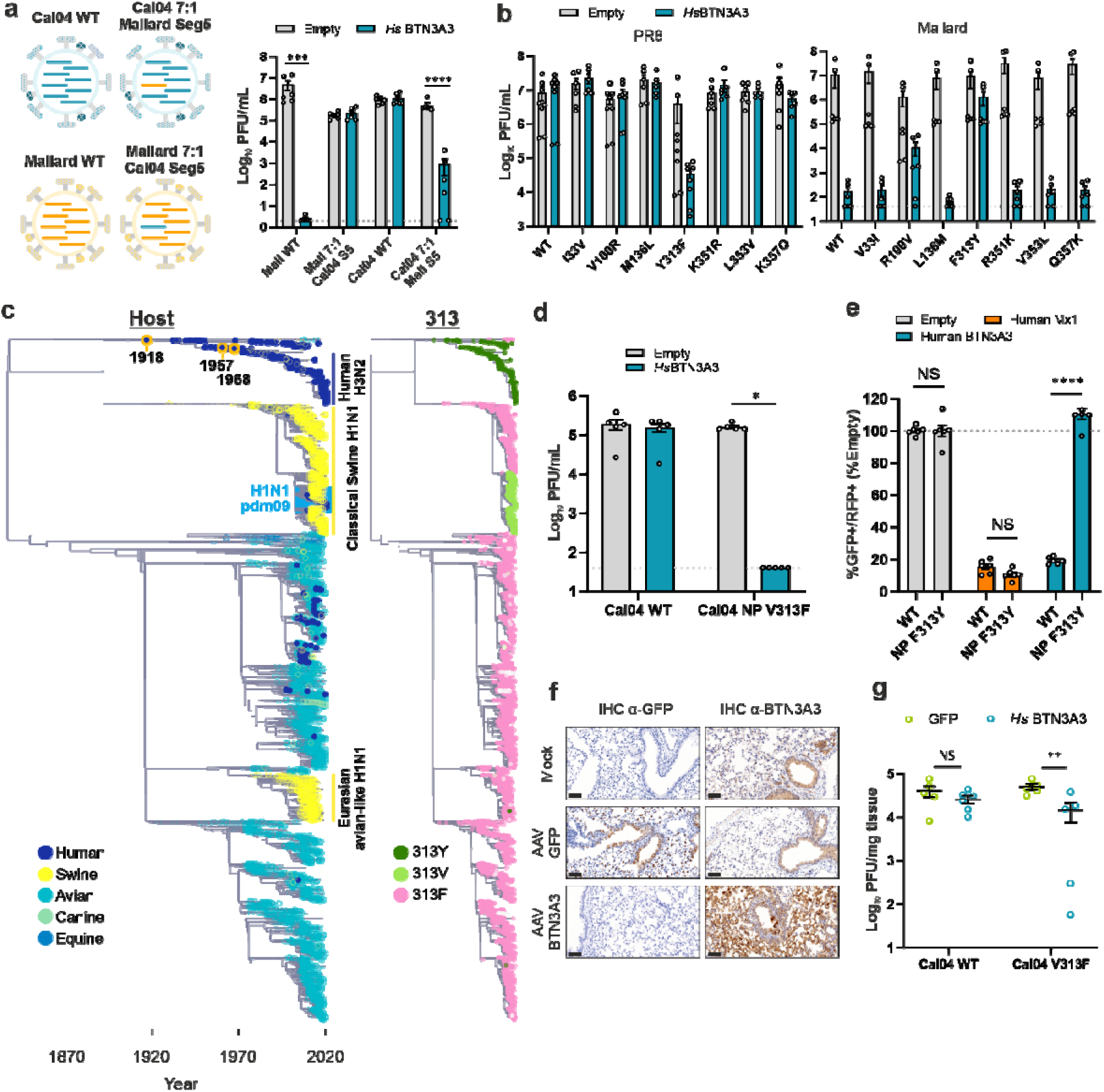
NP amino acid residue 313 determines sensitivity or resistance to BTN3A3. **a**, A549-Empty and BTN3A3 expressing cells were infected with the four viruses represented in the schematic at an MOI of 0.001 for 48 h, after which infectious virus titres were quantified by plaque assays. Data are mean +/- SEM of 3 independent experiments. Dots represent individual technical replicates. Statistical significance between groups was measured by multiple t-tests and corrected for multiple comparisons using the Holm-Šídák method. NS-Non-significant, *p≤0.05, **p≤0.01, ***p≤0.001, ****p≤0.0001. **b**, PR8 and Mallard NP single mutants were used to infect A549-Empty and A549-BTN3A3 and infectious titres were assessed by plaque assays as in (a). **c**, Tip-dated maximum likelihood phylogeny of the filtered IAV NP coding sequence dataset (see Methods). Tip shapes are annotated by host (left) and 313 residue (right, only residues occurring in more than 5% of the sequences are shown in the legend). Key clades and pandemic isolates are highlighted on the phylogeny. **d**, Cal04 and point mutant Cal04 V313F were used to infect A549-Empty and A549-BTN3A3 at a MOI of 0.001 for 48 hours, after which infectious virus titres were quantified by plaque assays and in (a). Data are mean +/- SEM of 2 technical replicates from 3 independent experiments. Statistical differences between Empty and *Hs*BTN3A3 were calculated using multiple t-tests and corrected for multiple comparisons using the Holm-Šídák method. NS-Non-significant, *p≤0.05. **e**, A549 overexpressing human Mx1 and BTN3A3 from SCRPSY lentiviral vectors were infected with GFP-tagged Mallard WT and NP F313Y mutant. 10 h post-infection cells were trypsinised, fixed and RFP-and GFP-positive cells were measured by FACS. Data are mean +/- SEM of 2 technical replicates from 2 independent experiments. Statistical differences between Empty and *Hs*BTN3A3 were calculated using multiple t-tests and corrected for multiple comparisons using the Holm-Šídák method. NS-Non-significant, ****p≤0.0001. **f**, Lungs from C57BL/6 mice instilled with PBS (mock) or AAV vectors containing either GFP or BTN3A3 were examined for expression of GFP or BTN3A3 after 3 weeks. Immunohistochemistry of mock-transduced-mice and AVV-GFP mice showed a weak BTN3A3 staining (light brown) in the bronchial epithelium, while AAV-*Hs*BTN3A3 overexpressing mice show a strong signal in pneumocytes, vessels and immune cells (dark brown). Bars represent 70μm. g, Mice were infected with either WT Cal04 (6:2 PR8 HANA) or Cal04-V313F (6:2 PR8 HANA) 3 weeks post-AAV treatment. Viral titres were assessed 3 days post infection. Data represent mean +/- SEM. Dots represent individual mice. Statistical differences between AAV-GFP and AAV-BTN3A3 were calculated using multiple t-tests and corrected for multiple comparisons using the Holm-Šídák method. NS-Non-significant, **p≤0.01.

We next reconstructed a time-calibrated phylogenetic tree for a comprehensive set of more than 30,000 IAV NP sequences (Fig. 4c). Human NP sequences have almost exclusively 313Y or 313V while all NP clades circulating in avian hosts, as well as the Eurasian avian-like H1N1 swine clade, are dominated by strains with the conserved 313F residue. Less than 1% of avian IAV NP sequences contain 313L, which also confers susceptibility to BTN3A3 (Extended Data Fig. 7). Occurrence of the BTN3A3-resistant 313Y is specific to a human NP clade, derived from the H1N1 1918 pandemic, subsequently reassorted into the 1957 and 1968 pandemics and currently circulating as seasonal H3N2. Precise dating of the original F313Y change is difficult due to the small number of 1918 IAV genomes available. However, recently sequenced early pandemic genomes all have a 313Y in NP^35^, suggesting that F313Y took place prior to or soon after human emergence of the 1918 H1N1 human strain. 313V is specific to the classical swine H1N1 NP subclade, which entered the human population in the 2009 IAV pandemic. BEAST (Bayesian Evolutionary Analysis Sampling Tree) tip-dating analysis supports a F313V change taking place in swine hosts between mid-2002 and the end of 2006 (Text S1, Extended Data Fig. 8). Interestingly, the NP of the H1N1 2009 pandemic virus, was acquired from this 313V subclade. We have therefore tested if, similarly to PR8, the reversion of NP V313F in a pmd09 virus would result in a BTN3A3 sensitive virus. Indeed, unlike its wild-type (WT) counterpart, the Cal04 NP V313F mutant showed a very pronounced restriction in BTN3A3 overexpressing cells (Fig. 4d). Interestingly, residue 313 had been previously associated with evasion of Mx1 resistance^19,20^. Avian-to-human mutation F313Y, when combined with R100V and/or L283P, exhibits increased Mx1 sensitivity^19^. However, here we observed that the single F313Y mutation while it is sufficient to overcome BTN3A3, does not overcome Mx-1 restriction (Fig. 4e). In addition, we showed in loss of function experiments that BTN3A3 restriction occurs in an Mx1-independent manner (Extended Data Fig. 9).

We further validated the role of NP residue 313 as a determinant of resistance to BTN3A3 in an experimental model of influenza *in vivo*. B6 mice were intranasally transduced with AAV6.2FF vectors expressing either BTN3A3 or GFP and infected with a reassortant Cal04 (6:2 PR8 HA/NA) containing the glycoproteins of PR8 to maximise replication in mouse tissues, with or without the V313F mutation. As shown in similar studies^36^, the AAV achieved efficient expression of our genes of interest in the mouse respiratory tract. We observed abundant GFP (control) and BTN3A3 expression in the respiratory tract of mice at 3 weeks post AAV-transduction (Fig. 4f). In challenged mice, Cal04 (6:2 PR8 HA/NA) virus replication in the lungs was not significantly impaired in BTN3A3 transduced mice compared to controls. Conversely, titres reached by the single mutant Cal04-V313F in BTN3A3 transduced mice were significantly lower than those reached in GFP transduced mice (Fig. 4g), indicating that BTN3A3 restriction can also occur *in vivo*.

### BTN3A3 interferes with avian IAV viral RNA transcription

NP is an essential protein for the life cycle of IAVs. Its primary purpose being the encapsidation of the viral genome to ensure effective RNA transcription, replication and packaging^37^. As well as being a structural RNA-binding protein, NP is also a key adapter between virus and host cell processes as it is essential for the nucleo/cytoplasmic trafficking of viral ribonucleoproteins (vRNPs)^37–39^. Hence, we next tested whether BTN3A3 inhibits vRNP nuclear import by performing synchronised infections of A549-empty or A549-BTN3A3 followed by nuclear/cytoplasmic fractionation at early time points. At 45 minutes post infection, levels of Mallard vRNP-containing proteins in both cytoplasmic and nuclear fractions were equal between A549-Empty and BTN3A3 expressing cells, suggesting equally efficient virus entry and vRNP nuclear import (Fig. 5a left panel, and related quantification in Extended Data Fig. 10). At 90 minutes post infection, by which point the transcription and translation of viral genes had resulted in an increase in viral protein, a difference in viral protein levels was seen between the two cell lines in both cytoplasmic and nuclear fractions (Fig. 5a middle panel, Extended Data Fig. 10). This difference was further amplified at 6 hpi (Fig. 5a right panel, Extended Data Fig. 10). Hence, these data suggested that the initial stages of the virus life cycle including binding and entry, and likely vRNP nuclear import were largely unaffected by BTN3A3 overexpression while subsequent steps, such as viral transcription and/or translation were affected. To investigate this further, we next studied viral transcription and translation using minireplicon reporter assays. Using either vRNA- or cRNA-like reported plasmids, replication/transcription by full avian or 313F recoded mammalian RNPs were significantly reduced in BTN3A3 overexpressing cells (Fig. 5b; Extended Data Fig. 11). In principle, this could be due to effects on viral transcription, translation or protein stability. In infected cells, viral transcription activity can be analysed by measuring vRNA, mRNA and cRNA levels through time. When infecting cells with Mallard WT (Fig. 5c, top row), initial vRNA levels were equal between A549-Empty and A549-BTN3A3 cells. However, a 7.5-fold difference in mRNA levels was observed from as early as 30 minutes post infection in BTN3A3 expressing cells. This was matched by a similar reduction in the accumulation of cRNA and vRNA, which rely on the expression of new viral proteins. However, when using the BTN3A3-resistant mutant Mallard NP F313Y, there were no differences in vRNA, mRNA and cRNA levels in the presence or absence of BTN3A3 (Fig. 5c, bottom row). We concluded that BTN3A3 can restrict sensitive IAVs at the level of viral transcription.

**Figure 5:**
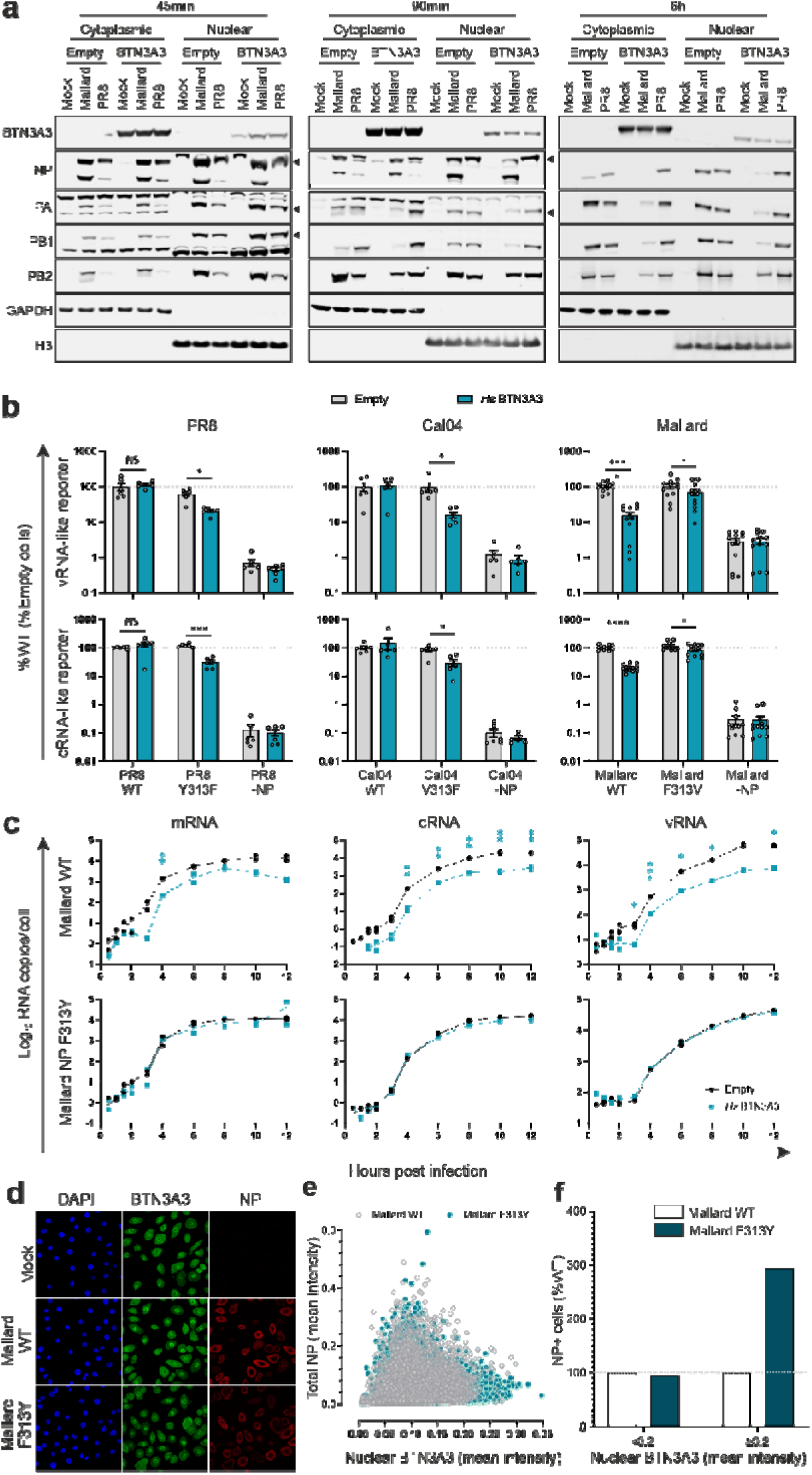
BTN3A3 blocks vRNA transcription. **a**, A549-Empty and A549-BTN3A3 cells were infected with PR8 and Mallard WT at an MOI 3 and cells were lysed at the indicated times post-infection. Nuclear/cytoplasmic fractionation was performed and viral RNP components were detected in either fraction by western blotting. GAPDH and Histone H3 were used as cytoplasmic and nuclear marker, respectively. Given the different times post infection and the different amounts of accumulated viral proteins between them, the three presented data sets originated from different volumes of lysate and different exposure times. **b**, A549-Empty and A549-BTN3A3 cells were transfected with pDUAL plasmids encoding for PB2, PB1, PA and NP of the indicated viruses alongside firefly luciferase-coding vRNA- or cRNA-like reporter plasmids. 48 h post transduction, cells were lysed and firefly luciferase was measured. Values were normalised to WT NP-coding RNPs transfected in A549-Empty. Data are mean +/- SEM of 2 technical replicates from 3 independent experiments. Statistical differences between Empty and *Hs*BTN3A3 were calculated using multiple t-tests and corrected for multiple comparisons using the Holm-Šídák method. NS= non-significant, *p≤0.05, **p≤0.01, ***p≤0.001, ****p≤0.0001. **c**, A549-Empty and A549-BTN3A3 cells were infected at MOI 3 and cells were lysed at the indicated times post infection. Total RNA was extracted, and different viral RNA species were quantified by a 2-step RT-qPCR. Data are mean +/- SEM of 3 independent experiments. Statistical differences between Empty and *Hs*BTN3A3 for each time point were calculated using multiple t-tests and corrected for multiple comparisons using the Holm-Šídák method. *p≤0.05, **p≤0.01, ***p≤0.001. **d**, Representative image of confocal microscopy of hBEC3-KT cells infected with Mallard WT or Mallard NP F313Y at MOI 3. 6 hours post infection, cells were immunostained with NP (red) and BTN3A3 (green). DAPI staining was used as nuclear marker. Image acquisition was performed with a Zeiss 880 confocal microscope using a 63x objective. **e**, Images from >3500 cells from four independent experiments performed as in d were used to quantify total NP and nuclear BTN3A3 for Mallard WT and Mallard NP F313Y. **f**, Values from b were stratified based on nuclear BTN3A3 intensity (<0.2 or ≥0.2). Data represents relative abundance of total infected cells present in each of the two nuclear BTN3A3 intensities ranges, taking values obtained with Mallard WT as 100%.

Interestingly, we also found components of the ribonucleoprotein complex to co-immunoprecipitate with BTN3A3. However, to overcome the non-specific binding of NP to Protein A agarose beads, the BTN3A3 IP was performed using the supernatants of NP immunoprecipitates (Extended Data Figure 12). IAV transcription takes place in the nucleus, and despite BTN3A3 being a transmembrane protein, localisation of constitutively expressed BTN3A3 in hBEC3-KT cells was predominantly nuclear (Fig. 5d). Image analysis of infected cells showed that, when cells express high levels of nuclear BTN3A3, viral protein synthesis (measured by NP accumulation) of Mallard WT was weaker (Fig. 5d-f and Extended Data Fig. 13a-c). Additionally, NP accumulation from a BTN3A3-resistant mutant was less sensitive to high levels of nuclear BTN3A3. We obtained similar data using the same assays with Cal04 (BTN3A3 resistant) and the BTN3A3-sensitive mutant Cal04 NP V313F (Extended Data Figure 13).

### NP residue 52 is a key determinant of sensitivity/resistance to BTN3A3 associated with recent avian IAVs spillovers into humans

We next analysed whether BTN3A3 evasion could correlated with the zoonotic potential of avian IAVs. We focused on H7N9 avian IAVs whose NP contains a 313F residue and theoretically should be restricted by BTN3A3. However, their high number of human infections^40^ led us to hypothesise that they might overcome BTN3A3 restriction, through an alternative mechanism. We therefore used a 6:2 H7N9 reassortant (containing the glycoproteins from PR8 and the internal segments from H7N9) and infected A549-Empty and A549-BTN3A3. The reassortant H7N9 was not restricted significantly by BTN3A3, despite possessing 313F in NP, whereas as a 5:3 H7N9 reassortant containing Mallard NP was restricted ∼70 fold on average (Fig. 6A). Comparison of H7N9 and Mallard NP sequences highlighted differences at 8 residue positions (34, 52, 186, 352, 373, 377, 406 and 482). To determine whether any of these amino acid substitutions could enable H7N9 viruses to evade BTN3A3 restriction, we constructed and tested single residue substitution mutants, and identified that the NP substitution N52Y rendered H7N9 sensitive to BTN3A3 (Fig. 6a). The effect of 52N was only seen in the presence of 313F; the N52Y mutation was not sufficient to confer BTN3A3-sensitivity in the presence of 313Y (Figure 6b). Unlike NP residue 313, there are no distinct host-clade associations with the identity of amino acid residue at position 52 (Fig. 6c). Instead, 52N-coding NP sequences occur in multiple independent avian IAV lineages. We noted that, with the exception of the H5 serotype, position 52 of human isolates from the avian clade have not only residue N but also residues Q or H (Fig. 6d). We show that mutants engineered with the 52Q and 52H in either the Mallard or the H7N9 backgrounds result in BTN3A3 evasion as efficiently as those possessing 52N (Fig. 6e), further reinforcing the hypothesis that BTN3A3 evasion is a key phenotype of avian IAVs spilling over into humans. Interestingly, despite being separated in the primary sequence of NP, positions 52 and 313 are closely juxtaposed within the NP head domain (Fig. 6f). Moreover, both residues are located on the surface of the NP trimer, indicating that they would be accessible to interactions with other viral and host factors when NP oligomerises to form a vRNP (Figure 6g). Of note, 52Y exhibits a protuberant alkaline methylphenol-containing side chain, while N, Q and H side chains are polar, likely not occupying the same space around 313F (Fig. 6h).

**Figure 6:**
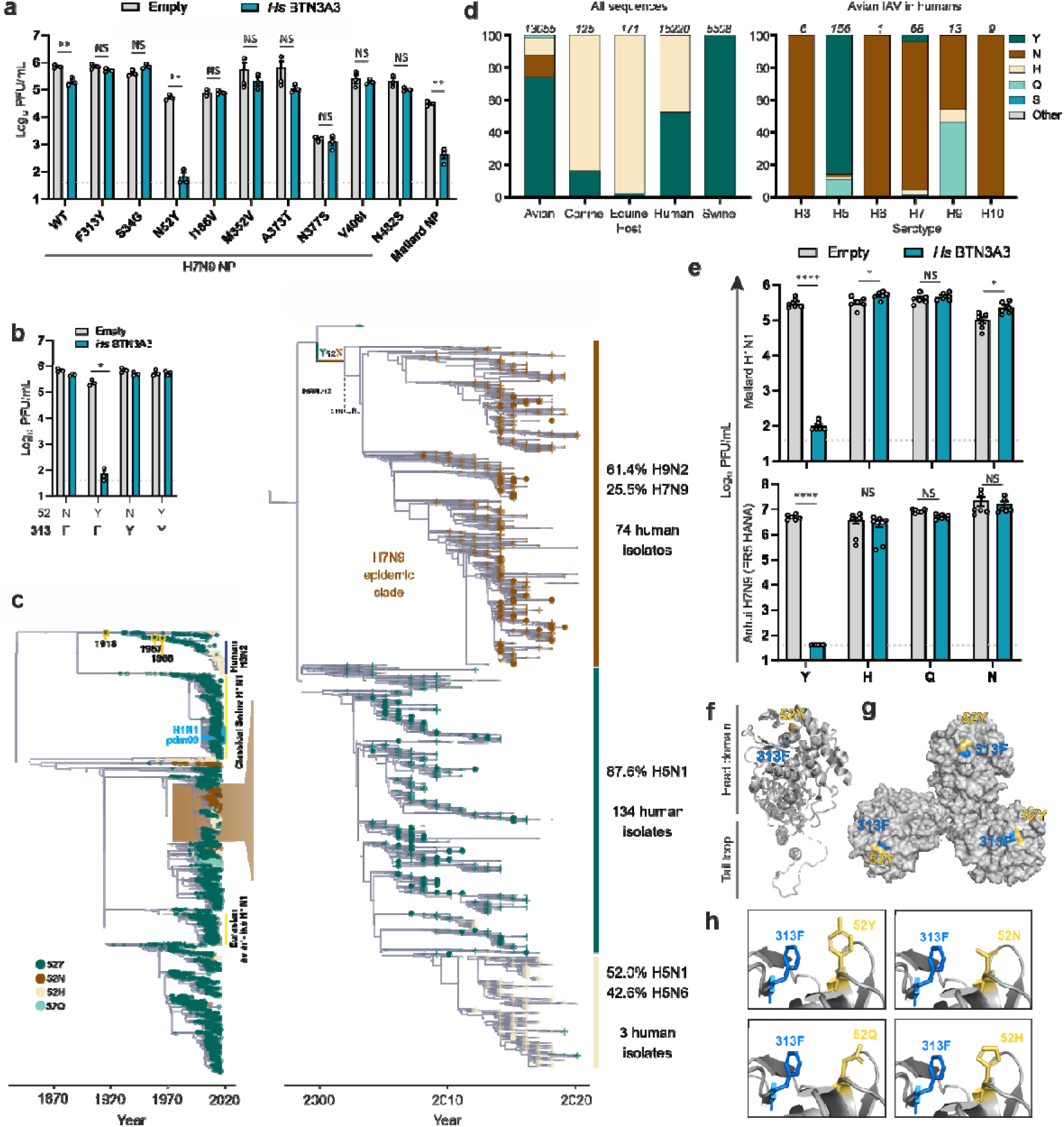
Evasion of BTN3A3 correlates with avian-to-human spillovers. **a-b**, Replication of H7N9:2 PR8 HA/NA reassortant viruses with the indicated NP mutations in A549-empty or A549-BTN3A3 expressing cells. Cells were infected at an MOI of 0.001 for 48h and titres were measured by plaque assay. Evasion of BTN3A3 restriction maps to amino acid residue 52 (a) but it is dependent on residue 313F (b). Data are mean +/- SEM of 3 independent experiments. Data are mean +/- SEM of a single technical replicate from 3 independent experiments. Statistical differences between values obtained in empty and BTN3A3 expressing cells were calculated using multiple t-tests and corrected for multiple comparisons using the Holm-Šídák method. NS =non-significant, *p≤0.05, **p≤0.01, ***p≤0.001, ****p≤0.0001. **c**, Tip-dated maximum likelihood phylogenies of the filtered IAV NP coding sequence dataset (left) and of all sequences clustering within the highlighted highlighted avian IAV clade (right). Tip shapes are coloured by site 52 residues (only residues present in more than 1% of the sequences are shown in the legend). Right: human isolates are annotated with circles and non-human isolates with transparent crosses to highlight human transmission. The branch where the Y52N change took place is annotated on the tree. Statistics of serotypes (present in more than 10% of the sequences in each subclade) and the number of human isolates are shown on the right of each subclade. **d**, identity of amino acid residue in position 52 of all NP proteins available in the NCBI flu dataset used in this study (see Methods) (left) and in those collected from spillovers of avian IAV in humans (right). Sequences were separated into hosts and virus subtypes and occurrence of different 52 residues was measured. Numbers over the bars equate to the number of sequences available for that particular host/subtype. **e.** Infectious virus titres obtained in A549-Empty and BTN3A3 expressing cells with NP 52 mutants of either Mallard or H7N7 6:2 PR8. Cells were infected at an MOI of 0.001 for 48h. Infectious virus titres were acquired by plaque assay. Data are mean +/- SEM of 2 technical replicates from 3 independent experiments. Statistical differences between Empty and BTN3A3 expressing cells were calculated using multiple t-tests and corrected for multiple comparisons using the Holm-Šídák method. NS= non-significant, *p≤0.05, **p≤0.01, ***p≤0.001, ****p≤0.0001. **e**, Ribbon structure of the monomer form of NP with amino acid residues changes mapped in colour. Using Pymol software and the 3.2Å crystal structure (Protein Data Base ID 2IQH); NP positions 52 and 313 are highlighted in yellow and blue, respectively. **g**, Surface structure of trimeric NP with 52 and 313 residues visibly mapped. **h**, Location and side chains of 313F and different 52 mutations were modelled using Pymol. Four focused panels are 52Y, 52N, 52D and 52Q.

Interestingly, NP residues 52 has also been implicated in the evasion of Mx1 resistance^19,20^. However, Mx1 and BTN3A3 differ in how viral sensitivity/resistance depends on NP residues 52 and 313 (Extended Data Fig. 14). The evasion of BTN3A3 restriction conferred by residue 52, is dependent on residue 313, which is not the case for evasion of Mx1 restriction.

## Discussion

IAVs host range is determined by multiple species-specific interactions between viral and cellular factors^12^. Our data reveal that human BTN3A3 is a powerful barrier for avian, but not human IAV replication. BTN3A3 interferes with avian IAV viral RNA transcription. Butyrophilins were discovered originally as proteins involved in lactation and milk production, but they are now known to have immunomodulatory functions^3,40–42^. These group of proteins have not been generally associated with antiviral properties, although one of two studies using ISG libraries against Ebola virus replicons identified BTN3A3 as a restriction factor for this virus^43,44^.

IAVs susceptibility to BTN3A3 restriction is determined by the NP protein and maps primarily to amino acid residues 313 and 52. Avian viruses almost universally possess NP 313F (resulting in a BTN3A3 susceptible genotype). A small minority (less than 1%) of avian IAVs possess the 313L residue, which also confers susceptibility to BTN3A3. Human viruses almost always possess either 313Y or 313V (BTN3A3 resistant genotype).

The anti-IAV properties of BTN3A3 arose in primates. Hence, humans are the only species naturally infected with IAVs that restrict avian IAVs replication through this effector. Neither avian species nor swine, dogs or horses impose a BTN3A3-related selective pressure on the NP 313 residue. Importantly, the swine 2009 pandemic originated from a BTN3A3 resistant NP 313V clade, that we estimate appeared in pigs between 2002 and 2006, ahead of the 2009 pandemic (Extended Data Fig. 8). The strong conservation of residue 313F in avian IAVs is suggestive of purifying selection on this site during circulation in birds. Conversely, strains with both 313F and V genotypes circulate in pigs. This indicates relaxed selection on the virus for 313F in swine, potentially allowing for occasional changes to BTN3A3-resistant residues. While the current data support a F313V change during circulation of the virus in swine ahead of the 2009 pandemic, there are no sequences available of the earliest intermediate viruses in this clade. Given the reassortment promiscuity of these viruses and reports of sporadic swine-to-human transmission in the late 1990s and early 2000s^45^, the residue change taking place in a non-swine host followed by a back-spillover event remains a possibility for the origin of the 2009 pandemic virus.

We show that evasion of BTN3A3 restriction is a conserved trait of avian viruses that have successfully spilled over into humans. To date, avian influenza viruses of subtypes H3, H5, H6, H7, H9 and H10, have crossed the species barrier and caused human infections^9^, despite maintaining the avian-like NP-313F. We have shown that viruses that evade BTN3A3 restriction have mutated from a BTN3A3-sensitive 52Y residue to a BTN3A3-resistant N, Q or H residue. We hypothesise that residue 52 masks the effect of 313F. Interestingly, the majority of IAV isolates from horses and dogs also contains a resistant BTN3A3 genotype (313F and 52H). Similar examples of “avian-to-human” adaptive mutations have occurred in PB2, most notably at residue 627^12,46^ which is a glutamic acid in avian viruses and adapts to a lysine in human viruses such as those that caused the 1918, 1957 and 1968 pandemics, and their seasonal descendant viruses. However, PB2 adaptation of the 2009 pandemic virus occurred through alternative mutations in amino acid residues 271, 590 and 591, which are structurally close to 627^47^.

An apparent exception to BTN3A3 representing a key barrier to avian IAVs spillover into humans are avian H5 viruses, which can infect humans despite many of them possessing a BTN3A3-susceptible genotype (52Y, 313F). The viruses of the H5 subtype are highly pathogenic strains containing polybasic cleavage sites that may allow a wider cellular tropism, and potentially infect low-BTN3A3 expressing cells, leading to a higher chance of a successful infection^48,49^. Alternatively, other compensatory mutations, even in segments other than segment 5, may provide resistance. In recent years, the A/Goose/Guangdong/1/96 (GsGd) lineage of highly pathogenic avian influenza (HPAI) H5 virus, originally emerged in Asia, have caused an increasing number of outbreaks in wild birds and poultry across multiple continents^50^. The frequency of these outbreak has increased substantially in Europe since late 2020. Worryingly, out of 480 avian H5N1 isolates sampled between January 2021 and June 2022 in Europe (Tables S3-S4), 277 have a BTN3A3 resistant genotype^50^. In the same period, transmission of HPAI H5 viruses to humans, and other wild mammals, have been reported both in Europe, Asia and North America. 3 of 6 zoonotic cases have been caused by Gs/GD HPAI H5 viruses with a BTN3A3 resistant genotype (Table S5)^51^, providing opportunities for further virus adaptation to humans.

We noted how both NP residues 313 and 52 have been associated with evasion of Mx1 resistance^19,20^. We presented data *in vitro* clearly suggesting that despite affecting similar stages of the virus life cycle, and through common amino acid residues, BTN3A3 acts independently from Mx1. Also, our experiments *in vivo* were carried out in regular C57BL/6 mice transduced with AAV vectors expressing BTN3A3. C57BL/6, like many other laboratory mouse strains possess a non-functional Mx1, further reinforcing the point that BTN3A3 and Mx1 act independently from each other^27^. It is however relevant that two independent human restriction factors target a conserved surface of NP, suggesting that this constrained region is a key determinant of the zoonotic potential of avian IAVs.

Global efforts during the SARS-CoV-2 pandemic and over the years for IAV infections^2^ have shown that surveillance based on virus genomic sequences can be a useful tool not only to provide insight into disease spread and epidemiology, but also for the early identification of viruses with specific undesirable phenotypic traits from a public health perspective. With the possible exception of some H5 sub-types, a BTN3A3 resistant genotype appears to be a pre-requisite for avian IAV spillover in humans. Among the ∼13,000 avian IAV NP sequences available in the present dataset, we found 151 independent avian NP lineages which already apparently possess a BTN3A3 resistant genotype. Members of these lineages have been sampled across the globe, with hotspots in China and North America correlating with sampling efforts (Extended Data Fig. 15). Hence, it is likely unpredictable where the next BTN3A3 resistant lineage will occur. Overall, the data presented in this study provides a new outlook on the zoonotic potential of avian IAVs. Identifying the viral genotypes that determine BTN3A3 resistance/evasion can be a powerful additional marker for assessing the zoonotic potential of avian and animal IAVs and should be considered in risk-assessment frameworks^2^.

## Materials and methods

### Cells and viruses

MT4 suspension cells were cultured in RPMI (RPMI, Gibco) 1640 medium supplemented with 10% foetal bovine serum (FBS, Gibco), 100 U/mL penicillin and 100 µg/mL streptomycin. Madin-Darby canine kidney (MDCK) cells (ATCC), human embryonic kidney cells (293T) (ATCC), human adenocarcinomic alveolar basal epithelial cells (A549) (ATCC) were cultured in Dulbecco’s modified Eagle’s medium (DMEM, Gibco) supplemented with 10% foetal bovine serum (FBS, Gibco), 100 U/mL penicillin and 100µg/mL streptomycin. MDCK cells expressing Sialyltransferase 1 (MDCK-SIAT, kindly gifted by Prof John McCauley, The Francis Crick Institute) were further supplemented with 50 µg/mL geneticin.

hTERT-immortalized primary human foetal lung fibroblasts were cultured in minimum essential medium supplemented with 10% FBS, 100 U/mL penicillin, 100 µg/mL streptomycin and 1x non-essential amino acids. Normal human bronchial epithelial cells immortalised with CDK4 and hTERT (hBEC3-KT)^52^ were kept in Keratinocyte serum-free medium (Gibco) supplemented with 50 µg/mL of bovine pituitary extract, 5 ng/mL of human recombinant epidermal growth factor, 100 U/mL penicillin and 100 µg/mL streptomycin.

Culture media of SCRPSY-modified cells was further supplemented with 1 µg/mL of puromycin dihydrochloride. All cell lines were regularly tested for mycoplasma contamination and passaged once or twice weekly.

PR8, Cal04, Mallard and Anhui H7N9 viruses were generated by reverse genetics essentially as described previously^53^. Briefly, ∼2×10^6^ 293T cells were transfected in OptiMEM with pDUAL reverse genetics plasmids (250 ng plasmid for each virus segment), using 4uL of Lipofectamine 2000 according to manufacturer’s instructions. Twenty-four hours post transfection, the media was changed to serum-free DMEM supplemented with 0.14 % Bovine Serum Albumin (w/v) and 1 μg/mL L-(tosylamido-2-phenyl) ethyl chloromethyl ketone (TPCK)-treated trypsin. Virus-containing supernatant was collected after a further 2-day incubation and propagated in MDCK cells. Clarified supernatant was collected when approximately 90% of cells demonstrated cytopathic effect (typically 36 – 72 h post-infection).

### Arrayed ISG overexpression screening

Lentiviral transduction/flow cytometry-based screening was performed as previously described^24,25^. Briefly, MT4 cells were seeded in 96-well plates and further transduced with two SCRPSY-based lentiviral vector libraries consisting of 527 human and 344 macaque ISGs. At 48 hours post transduction, cells were infected with either PR8-GFP, Mallard-GFP or Cal04 7:1 PR8 segment 8 GFP (rescued as previously described^54^). After 10 h, cells were fixed and the percentage of transduced (TagRFP-positive) and IAV-infected (GFP-positive) cells was determined by flow cytometry using a Guava EasyCyte flow cytometer (Millipore).

The screens in figure 3A were performed in a similar manner. MT4, AA2 A549 or A549 overexpressing human ACE2 were transduced with lentiviral ISG-expressing library as described above^55^. Forty-eight hours post transduction, cells were infected with reporter-expressing virus aiming to achieve a 10-50% infection. The panel of viruses used in figure 3A was created with the following viruses: AdV, Human mastadenovirus C (Adenovirus 5); PRV, Suid herpesvirus 1 (Pseudorabies); HSV-1, Human herpesvirus 1; BoHV-1, Bovine herpesvirus 1; RVFV, Rift Valley fever phlebovirus; SFTSV, Dabie bandavius (Severe fever with thrombocytopenia syndrome virus); BUNV, Bunyamwera orthobunyavirus; BTV, Bluetongue virus; Rotavirus, Simian Rotavirus A/SA11; CHPV, Chandipura vesiculovirus; VSV, Indiana vesiculovirus; SFV, Semliki Forest virus; CHIKV, Chikungunya virus; HIV-1, Human immunodeficiency virus 1; IAV PR8, A/Puerto Rico/8/1934 (H1N1); IAV Cal04, A/California/04-061-MA/2009 (H1N1) rescued with PR8 segment 8; IAV Mallard, A/Mallard/Netherlands/10-Cam/1999 (H1N1); MeV, Measles Ed-Zag vac; SeV, Murine respirovirus (Sendai virus); PIV5, Mammalian orthorubulavirus 5 (Parainfluenza virus 5 or simian virus 5); hMPV, human Metapneumovirus; hRSV, human orthopneumovirus (human respiratory syncytial virus); bRSV, Bovine orthopneumovirus (bovine respiratory syncytial virus); PIV3, human respirovirus 3 (Parainfluenza virus 3); and ZIKA, Zika virus.

### Influenza A virus infection

Monolayers of A549, MRC5T and hBEC3-KT cells were washed once with PBS and infected with virus diluted in serum-free medium for 1h at 35°C or 37°C (MOI used is specified in the relevant figures). Medium was replaced with complete maintaining medium (for qRT-PCR or western blot analysis) or serum-free medium supplemented with 0.14%BSA and 1 µg/mL TPCK-treated bovine pancreas trypsin (growth kinetics and quantification of infectious virus titres). Infectious titres were determined by plaque assay.

### Virus quantification by plaque assay

Confluent monolayers of MDCK cells (5×10^5^ or 1×10^6^ cells seeded the day before infection in 12- or 6-well plates, respectively) were washed once with PBS, infected with 10-fold serial dilutions of virus and incubated for 1 h at 37°C to allow virus adsorption to the cells. After virus removal, cells were overlaid with DMEM including 0.14% BSA, 1 µg/mL TPCK-treated trypsin and 1.2% Avicel and incubated for 3 days at 37°C, 5% CO2. After removing the overlay, cells were fixed with PBS/4%formaldehyde and stained with a 20%methanol/ 10% acetic acid/ 0.2% Coomassie Blue solution for at least 1h. Staining solution was rinsed under tap water, plates were air-dried and plaques were then counted. Plaque assays for the avian viruses A/wild-duck/Italy/17VIR6926-1/2017 H5N2, A/duck/Italy/18VIR4932-2/2018 H7N7 and A/turkey/Italy/16VIR8643-54/2016 H9N2 were performed as described but using MDCK cells overexpressing galline ANP32A. Plaque assays for Cal04 backbone viruses were performed essentially as described but using MDCK-SIAT cells and a 3-day incubation period at 35°C. Cells were then fixed, fixing solution was removed and cells were washed with PBS and permeabilised using PBS/0.2% (v/v) Triton X-100 for 10 minutes. After two PBS washes, cells were incubated with a mouse anti-NP antibody (diluted 1:1000 in PBS/ 2% BSA) for 1 h on a rocking platform at room temperature. Following two PBS washes, cells were incubated with goat anti-mouse IgG-HRP conjugated secondary antibody (diluted 1:1000 in PBS/ 2% BSA) for 1h in the same conditions. Cells were washed three times with PBS, TrueBlue peroxidase substrate was then added, and cells were incubated at room-temperature covered from direct light. When blue-stained plaques were visible, cells were washed with water, allowed to dry, and plaques were counted (under a stereo microscope, if required).

### siRNA-mediated gene silencing

MCR5T (5×10^5^ cells/well), hBEC3-KT (5×10^5^ cells/we//) or A549-based cells (3.5 ×10^5^ cells/well) were seeded before transfection of siRNA molecules targeting BTN3A3 or Mx1. When transfecting, 20 pmol of siRNA was used with 3 μL of DharmaFECT 2 (Horizon Discovery) per well according to the manufacturer’s protocol.

### Minireplicon polymerase reporter assay

Sub-confluent monolayers of A549-Empty and A549-HsBTN3A3 cells (2×10^5^ cells seeded in 24-well plates the previous day) were co-transfected in triplicate with 150 ng of each pDUAL plasmid encoding PB2, PB1, PA and NP along with 20 ng of transfection control plasmid (CMV-driven expression of Renilla luciferase) and 50ng of a PolI-driven expression of vRNA or cRNA-like firefly luciferase reporter plasmids. As a negative control, transfections lacking the NP plasmid (empty pDUAL vector was used to balance plasmid intake) were also performed. Two days post transfection, medium was removed and cells were lysed with 120 µL of reporter lysis buffer. Cell debris was scraped off, lysates were collected into clean tubes and clarified by centrifugation (10,000 rpm for 5 minutes at 4°C) in a benchtop centrifuge. Luminescence was measured from 50 μL of lysate in opaque 96-well plates by using the Dual-Luciferase® reporter assay system (injection speed: 200μL/second; gap: 0.5 seconds; integration time: 10 seconds).

### RT-qPCR

Total RNA of infected cells was extracted from culture supernatants using the RNAdvance Blood kit (Beckman Coulter Life Sciences), including a DNase treatment, following the manufacturer’s instructions. Extracted and DNase-treated RNA was used to perform strand-specific 2-step qRT-PCR targeting the three different influenza RNA species, as previously described^56^. Briefly, a 5’ tagged primer is used for the cDNA synthesis, the reverse complement of which was then used as a forward primer in the real-time PCR step. Therefore, even in the event of primer-unspecific cDNA synthesis only the cDNA generated with the tagged primer will be detected and amplified in the real-time PCR. cDNA synthesis was performed with 5 µL of extracted RNA using the RevertAid kit (Thermofisher) in a VeritiTM 96 well Thermal Cycler. qPCR reactions were performed with 2 µL of cDNA using the Brilliant III Ultra-Fast qPCR mastermix. qPCR was performed and analysed using the QuantStudio® 3 thermocycler and software systems (ThermoFisher Scientific). A list of primers and probes can be found in supplementary information. In vitro-synthesised T7-transcribed viral mRNA, vRNA and cRNA-like transcripts were diluted and used as a standard curve. Relative quantification of viral RNA species was performed against GAPDH.

### Cell fractionation

A549-Empty and A549-HsBTN3A3 cells were infected with PR8 and Mallard WT viruses at an MOI of 3 for either 45 minutes, 90 minutes or 6 h. Cells were then washed in ice-cold PBS and lysed with 50 mM Tris-HCl pH 7.5, 10 mM KCl, 5 mM MgCl2, 0.5% NP40, 1 mM DTT, 10mM sodium b-glycerophosphate and Halt Protease inhibitor (1X, EDTA-free) (Thermo Fisher, 78429). Lysates were kept on ice for ten minutes (gently vortexed every 2-3 minutes) and centrifuged at 1000g for 5 minutes at 4°C. The supernatant (cytoplasmic fraction) was transferred to a clean 1.5 mL tube and the nuclear fraction was washed thrice with the above-mentioned lysis buffer (centrifugations done for 5 minutes at 1000 g). The nuclear fraction was then incubated on ice for 30 minutes with 2X LDS (Thermo Fisher, NP0007), 2X sample reducing agent (Thermo Fisher, NP0004), Denarase (final activity – 25U/ml) (c-LEcta, 20804-100k) and 10 mM MgCl2. The nuclear fraction was sonicated ten times for 20 seconds with 10 second intervals, before being centrifuged at 12000 g for 15 minutes. The supernatant was collected, and the pellet discarded. The cytoplasmic fraction was denatured using 1X LDS and 1X sample reducing agent.

### Immunoprecipitation

hBEC3-KT cells were infected with Mallard WT and NP F313Y at an MOI of 3 for 6h and lysed as previously described^57^. Lysates were pre-clarified overnight at 4°C with protein A agarose beads (Sigma Aldrich, P3476). 4 mg of protein lysates were then incubated with 12 µg of either anti-NP or IgG control antibody overnight at 4°C. Supernatants from the α-NP and IgG control were further incubated overnight at 4°C with 12 µg of anti-BTN3A3 and IgG control antibody, respectively. Fifteen µL of Protein-A agarose beads were added for an hour the next morning. IPs were then washed with 50 mM Tris-HCl, 500mM sodium chloride and 0.5% TritonX-100 followed by 50 mM Tris-HCl only before being eluted using 2X LDS and 2X sample reducing agent. Immunoprecipitates were lysed and further subjected to immunoblotting targeting NP, PB2, PB1, PA and BTN3A3.

### Immunoblotting

Total cell, immunoprecipitates or lysates from fractionation experiments were heated at 80°C for 5 minutes and subjected to polyacrylamide gel electrophoresis using 3-(N-morpholino) propanesulfonic acid (MOPS) running buffer and transferred to polyvinylidene difluoride (PVDF) (Merck Millipore, IPFL00010) membranes at 30V for 90 minutes. Membranes were blocked for 1h with 1X Tris-buffered saline with 0.2% Tween® 20 (TBST)/5% milk, washed 4 times with 1xTBST and stained with primary antibodies diluted in 5% BSA BSA/0.01% sodium azide overnight at 4°C. After four 10-minute 1xTBST washes, secondary antibodies were diluted in blocking buffer and incubated for 1h at room temperature protected from direct light. Following four more washes in 1xTBST, membranes were imaged using the LI-COR CLx-Odyssey Imaging platform. Densitometry analysis was performed using the Image Studio™ Lite Software.

### Immunofluorescence staining

The desired cell lines were seeded on 13 mm round glass coverslips in 24 well dishes. After infection/transfection, cells were washed twice with PBS and fixed with 4% formaldehyde in PBS for 20 minutes followed by three PBS washes. Cells were permeabilised with PBS/1% Triton X-100 for 10 minutes at room temperature. Following three washes with PBS, cells were blocked with 1 mL PBS/1% BSA for 1 hour followed by incubation for 1 hour with 200 µL of primary antibodies at the appropriate dilutions (for the detection of BTN3A3 in hBEC3-KT cells, the primary antibody was incubated overnight at 4°C). Three 1 mL PBS/0.2% Tween20 washes were then executed to remove unbound antisera followed by incubation of secondary antibodies in the same volume. 4’,6-Diamidino-2-phenylindole (DAPI) (100 ng/mL) was diluted in blocking buffer and incubated for 10 minutes. When using HCS CellMask™ Deep Red Stain (2 µg/mL – ThermoFisher Scientific), this was added after DAPI staining for 30 minutes followed by three PBS washes. After labelling steps were completed, coverslips were washed three more times with water and mounted upside down on glass slides with AF1 Mounting media (CitiFluor^TM^). Coverslips were imaged using the ZEISS LSM 710 or LSM 880 (with Airyscan) confocal laser scanning microscopes. Quantification of nuclear and cytoplasmic proteins from confocal images was performed using a pipeline created on CellProfilerTM. Pipeline details are provided in supplementary information.

### Immunohistochemistry

Sections (3 μm thick) of formalin-fixed and paraffin-embedded (FFPE) human lung and respiratory nasal epithelium (female, 52 years old, Asian, post-mortem donor, normal tissue; code: AMS-41022-NE and 500029028-DS, Amsbio) were stained with a rabbit anti-BTN3A3 antibody (HPA 007904, Atlas antibodies) for 1.5 hours at room temperature after pressure cooking in sodium citrate. For visualisation, EnVision Detection System HRP, peroxidase/DAB (rabbit, K500711, Agilent) was used according to manufacturer’s instructions. The specificity of the antibody was validated on FFPE A549-Empty and A549-HsBTN3A3 cell pellets. Negative control sections included an isotype control on a consecutive serial section. Slides were scanned with a Leica Aperio Versa 8 slide scanner (Leica Biosystems) and images were acquired with a Aperio ImageScope software (Leica Biosystems).

Mouse lung tissues were stained with a rabbit anti-BTN3A3 antibody (HPA 007904, Atlas antibodies) and anti-GFP antibody (#2555S, Cell Signaling) for 1.5 hours at room temperature after pressure cooking in sodium citrate. For visualisation, EnVision Detection System HRP, peroxidase/DAB (rabbit, K500711, Agilent) was used according to manufacturer’s instructions.

### AAV vectors

AAV vector genome plasmids were engineered to encode a ubiquitous CASI promoter^58,59^ driving expression of the human BTN3A3 gene followed by a WPRE^59^ and a SV40 polyA signal all contained between AAV2 inverted terminal repeats. AAV vectors were produced by co-transfection of human embryonic kidney 293 cells with genome and packaging plasmids as described previously^59^. Vectors pseudotyped with AAV6.2FF were purified by use of a heparin column^59^. AAV vector titres were determined by quantitative polymerase chain reaction (qPCR) analysis as described elsewhere^60^.

### Mouse infections

Six week-old female C57BL/6 mice were purchased from Envigo (UK). Animals were maintained at the University of Glasgow under specific pathogen free conditions in accordance with UK home office regulations (Project License PP1902420) and approved by the local ethics committee. Following 7 days of acclimatization, the mice were briefly anesthetized using inhaled isoflurane and transduced with 1×10^11^ virus genomes of adeno-associated virus (AAV). Mice received either AAV-GFP or AAV BTN3A3 in 50 µL PBS intranasally. Twenty-one days post AAV instillation mice were infected with Cal04 and Cal04 V313F (both 6:2 with PR8 HA/NA). Mice were briefly anesthetized using inhaled isoflurane and intranasally infected with 5 ×10^3^ PFU of each virus in 20 μL of PBS. All mice were euthanized by cervical dislocation 3 days post infection. Weight loss for all groups did not exceed the humane cut-off point of 20%. Group sizes were based on previous experiments^61^. Once euthanised, mouse lungs were extracted and frozen at -80°C. Lungs were thawed, weighted and homogenised with 1 mL of DMEM. Homogenates were cleared by centrifugation (3000 rpm, 4°C) and titrated by plaque assay.

### *In silico* identification of BTN3 homologs

To identify proteins expressed by other species, homologous to the human BTN3 genes, we performed a Blastp search (v.2.8.1) against all available members of the Haplorrhini suborder in the NCBI blast protein refseq database version 5 (e value cutoff 1e-60, as of 6th of April 2021)^62^. The human BTN3A3 protein sequence (NP_008925.1) was used as a probe for the BLAST search. The isoform with the longest sequence was kept for each protein product annotated with the same name. Similarly, Blastp with human BTN3A3 was used for identifying proteins expressed by non-primate species susceptible to IAVs infection: *Gallus*, *Anas platyrhynchos*, *Equus caballus*, and *Sus scrofa* and more distant human paralogues (Table S1).

Protein members of the butyrophilin 3 subfamily retrieved from the BLAST search were manually cross-checked with proteins in the Ensembl database^63^ and if the protein sequences were not identical between the two databases the sequence with highest similarity to the human BTN3A3 protein sequence was retained. A total of 30 proteins were retrieved from the following species: Pan troglodytes, *Cebus imitator, Equus caballus, Homo sapiens, Gorilla gorilla gorilla, Chlorocebus sabaeus, Macaca mulatta, Pongo abelii, Carlito syrichta, Mandrillus leucophaeus, Callithrix jacchus, Nomascus leucogenys, Rhinopithecus roxellana*.

A custom set of Pfam hmm profiles were used for identifying the conserved domains in the proteins, comprising the immunoglobulin V-set domain (PF07686), CD80-like C2-set immunoglobulin domain (PF08205), PRY (PF13765) and SPRY (PF00622) domains. All protein sequences were scanned with the profile set using hmmscan (HMMER 3.3)^64^. The best hit for each identified domain was extracted from the protein sequence aligned with the respective domain segments using mafft (v7.453, --maxiterate 1000 --localpair)^65^. Protein alignments were converted to codon alignments using pal2nal^66^. Phylogenies for each separate domain alignment (Extended data Fig. 5) and concatenated domain sequences (Figure 3a) were reconstructed using iqtree (version 1.6.12)^67^ with the best suited substitution model selected by the iqtree ‘-m TEST’ option and 10,000 ultrafast bootstrap replicates.

#### IAVs phylogenetic analysis

A total of 35,477 full-length NP coding sequences unique on the nucleotide level (identical sequences collapsed) were retrieved from the NCBI Flu database (https://www.ncbi.nlm.nih.gov/genomes/FLU/Database/nph-select.cgi?go=database, as of the 8th of June 2021, sampled until the end of 2020), only including type A influenza sequences annotated to have been isolated from avian, canine, equine, human and swine hosts. Sequences with ambiguous nucleotides and internal stop codons were removed, resulting in a dataset of 34,079 sequences. The corresponding protein sequences were aligned using mafft (v7.453, --maxiterate 1000 --localpair)^65^ and then converted to a codon alignment with pal2nal^66^. Metadata associated with each sequence accession were retrieved and tabulated.

To reduce oversampling of related sequences the dataset was clustered with a minimum sequence identity of 0.99 using MMseqs2 (--min-seq-id 0.99 --cov-mode 0)^68^. One representative was kept from each cluster leading to a filtered dataset of 14,665 sequences. The codon alignment of the filtered set was used to reconstruct a phylogeny with iqtree under a GTR+I+G4 model (selected as the most appropriate model with the ‘-m TEST’ option)^67^. The resulting phylogeny was then time-calibrated using TreeTime^69^ (Fig. 4c). Eleven sequences with annotated dates inconsistent with the root-to-tip regression were subsequently excluded from the analysis.

To explore the H7N9 epidemic NP clade in more detail, representative sequences from this broader avian NP clade (Fig. 6c) as well as the unfiltered sequences from each representative’s corresponding cluster were retrieved (3,150 sequences). The codon alignment of these NP sequences was used to infer a more detailed maximum likelihood phylogenetic reconstruction of this particular clade (iqtree under a GTR+I+F+G4 model with 10,000 Ultrafast bootstrap replicates)^67,70^ and time-calibrated using TreeTime as described above^69^ (Fig. 6c). All phylogenies were visualized using the ggtree R package^71^, unless stated otherwise. Tree statistics were analysed using the ete3 Python package^72^.

### Data availability

Alignments and raw phylogenetic data related to this study can be found in the following GitHub repository: https://github.com/spyros-lytras/BTN3A3_IAV.

## Acknowledgments

We are thankful to the authors, originating and submitting laboratories of the sequences from GISAID’s EpiFlu™ Database on which some of this research is based (see Table S6). We thank Pablo Murcia (MRC-University of Glasgow Centre for Virus Research) for providing clinical virus isolates and fruitful discussions, and Professor Malik Peiris (The University of Hong Kong) who kindly gifted A/ruddy shelduck/Mongolia/963V/2009 (H3N8). We are grateful to Sushant Bhat, Munir Iqbal (The Pirbright Institute), Laurence Tiley, Ron Fouchier (Erasmus MC) and Daniel Perez (The University of Georgia) for providing the reverse genetics systems of the A/Anhui/1/2013 (H7N9) and A/Mallard/Netherlands/10-Cam/1999 (H1N1), A/Puerto Rico/8/1934 (H1N1) and A/California/04-061-MA/2009 (H1N1) viruses, respectively. We acknowledge Dr John Mccauley (The Crick Institute) for sharing the MDCK-SIAT cells. This work was supported by the UK Medical Research Council (MC_UU_12016/10) awarded to MP and SJW and the following grants: Wellcome Trust 206369/Z/17/Z (to MP); Biotechnology and Biological Sciences Research Council (BBSRC) BB/P013740/1 awarded to FG and PD; BBSRC BB/S00114X/1 awarded to FG, PD and SJW; EU Horizon2020: DELTA-FLU (no 727922) awarded to PD and IM; Natural Sciences and Engineering Research Council of Canada (NSERC) Discovery Grant (RGPIN-2018-04737) awarded to SKW; Daphne Jackson Fellowship funded by Medical Research Scotland awarded to SS; MRC Career Development Award and Transition Support Award (MR/N008618/1 and MR/V035789/1) to EH; Wellcome Trust 210703/Z/18/Z awarded to MKLM; Medical Research Council MC_UU_12014/5 awarded to CB. GQ and JH are funded by Medical Research Council MC_UU_12014/12.

## Author’s Contribution

<u>Conceptualization;</u> RMP, SJW, MP; <u>Methodology:</u> RMP, SB, MKZ, SL, SS, JCW, VH, KEH, LO, SKW, MKLM; <u>Software:</u> SL, JH, QG; <u>Validation:</u> RMP, SB, SL, MKZ, JCW, VH, KEH; <u>Formal analysis:</u> RMP, SB, SL, MKZ, SS, VH, MVk, NCR, MCR, MVa, JH; <u>Investigation:</u> RMP, SB, SL, MKZ, SS, JCW, VH, MVk, NCR, MCR, MVa, LO, AW, CL, YP, EV, MLT, WF, KEH, AT; <u>Resources:</u> RMP, AT, CB, EH, PD, IM, SKW, SJW, MP; <u>Data curation:</u> RMP, SL, MVk, NCR, MCR, MVa, AW, EV; <u>Writing-original draft preparation:</u> RMP, SB, SL, MKZ, JCW, VH, MLT, SKW, MP; <u>Writing-Review & editing:</u> RMP, SB, SL, MKZ, SS, JCW, VH, MVk, NCR, MCR, MVa, AW, CL, JH, EV, MLT, WF, KEH, QG, LO, AT, FG, EH, PD, IM, SKW, MKLM, SJW, MP; <u>Visualization:</u> RMP, SB, SL; <u>Supervision:</u> RMP, JH, CB, SKW, MKLM, SJW, MP; <u>Project administration:</u> SJW, MP; <u>Funding acquisition:</u> SS, CB, FG, PD, IM, SKW, MKLM, SJW, MP

## REFERENCES

1 Lipsitch, M. et al. Viral factors in influenza pandemic risk assessment. Elife 5, doi:10.7554/eLife.18491 (2016).

2 Burke, S. A. & Trock, S. C. Use of Influenza Risk Assessment Tool for Prepandemic Preparedness. Emerg Infect Dis 24, 471–477, doi:10.3201/eid2403.171852 (2018).

3 Afrache, H., Gouret, P., Ainouche, S., Pontarotti, P. & Olive, D. The butyrophilin (BTN) gene family: from milk fat to the regulation of the immune response. Immunogenetics 64, 781–794, doi:10.1007/s00251-012-0619-z (2012).

4 Ye, Q., Krug, R. M. & Tao, Y. J. The mechanism by which influenza A virus nucleoprotein forms oligomers and binds RNA. Nature 444, 1078–1082, doi:10.1038/nature05379 (2006).

5 Yoon, S.-W., Webby, R. J. & Webster, R. G. in Influenza Pathogenesis and Control - Volume I (eds Richard W. Compans & Michael B. A. Oldstone) 359–375 (Springer International Publishing, 2014).

6 Krammer, F., et al. Influenza. Nature Reviews Disease Primers 4, 3, doi:10.1038/s41572-018-0002-y (2018).

7 Harrington, W. N., Kackos, C. M. & Webby, R. J. The evolution and future of influenza pandemic preparedness. Experimental & Molecular Medicine 53, 737–749, doi:10.1038/s12276-021-00603-0 (2021).

8 Worobey, M., Han, G. Z. & Rambaut, A. Genesis and pathogenesis of the 1918 pandemic H1N1 influenza A virus. Proc Natl Acad Sci U S A 111, 8107–8112, doi:10.1073/pnas.1324197111 (2014).

9 Short, K. R. et al. One health, multiple challenges: The inter-species transmission of influenza A virus. One Health 1, 1–13, doi:10.1016/j.onehlt.2015.03.001 (2015).

10 Liu, D. et al. Origin and diversity of novel avian influenza A H7N9 viruses causing human infection: phylogenetic, structural, and coalescent analyses. Lancet 381, 1926–1932, doi:10.1016/s0140-6736(13)60938-1 (2013).

11 Wang, X. et al. Epidemiology of avian influenza A H7N9 virus in human beings across five epidemics in mainland China, 2013–17: an epidemiological study of laboratory-confirmed case series. The Lancet Infectious Diseases 17, 822–832, doi:https://doi.org/10.1016/S1473-3099(17)30323-7 (2017).

12 Long, J. S., Mistry, B., Haslam, S. M. & Barclay, W. S. Host and viral determinants of influenza A virus species specificity. Nature Reviews Microbiology 17, 67–81, doi:10.1038/s41579-018-0115-z (2019).

13 Rogers, G. N. & Paulson, J. C. Receptor determinants of human and animal influenza virus isolates: differences in receptor specificity of the H3 hemagglutinin based on species of origin. Virology 127, 361–373, doi:10.1016/0042-6822(83)90150-2 (1983).

14 Di Lella, S., Herrmann, A. & Mair, C. M. Modulation of the pH Stability of Influenza Virus Hemagglutinin: A Host Cell Adaptation Strategy. Biophys J 110, 2293–2301, doi:10.1016/j.bpj.2016.04.035 (2016).

15 Zaraket, H. et al. Increased acid stability of the hemagglutinin protein enhances H5N1 influenza virus growth in the upper respiratory tract but is insufficient for transmission in ferrets. J Virol 87, 9911–9922, doi:10.1128/jvi.01175-13 (2013).

16 Long, J. S. et al. Species difference in ANP32A underlies influenza A virus polymerase host restriction. Nature 529, 101–104, doi:10.1038/nature16474 (2016).

17 Blumenkrantz, D., Roberts, K. L., Shelton, H., Lycett, S. & Barclay, W. S. The short stalk length of highly pathogenic avian influenza H5N1 virus neuraminidase limits transmission of pandemic H1N1 virus in ferrets. J Virol 87, 10539–10551, doi:10.1128/jvi.00967-13 (2013).

18 Park, S. et al. Adaptive mutations of neuraminidase stalk truncation and deglycosylation confer enhanced pathogenicity of influenza A viruses. Sci Rep 7, 10928, doi:10.1038/s41598-017-11348-0 (2017).

19 Mänz, B. et al. Pandemic influenza A viruses escape from restriction by human MxA through adaptive mutations in the nucleoprotein. PLoS Pathog 9, e1003279, doi:10.1371/journal.ppat.1003279 (2013).

20 Riegger, D. et al. The nucleoprotein of newly emerged H7N9 influenza A virus harbors a unique motif conferring resistance to antiviral human MxA. J Virol 89, 2241–2252, doi:10.1128/jvi.02406-14 (2015).

21 Zhu, W. et al. A Gene Constellation in Avian Influenza A (H7N9) Viruses May Have Facilitated the Fifth Wave Outbreak in China. Cell Rep 23, 909–917, doi:10.1016/j.celrep.2018.03.081 (2018).

22 Chen, Y. et al. Rare variant MX1 alleles increase human susceptibility to zoonotic H7N9 influenza virus. Science 373, 918–922, doi:10.1126/science.abg5953 (2021).

23 Shaw, A. E. et al. Fundamental properties of the mammalian innate immune system revealed by multispecies comparison of type I interferon responses. PLoS Biol 15, e2004086, doi:10.1371/journal.pbio.2004086 (2017).

24 Kane, M. et al. Identification of Interferon-Stimulated Genes with Antiretroviral Activity. Cell Host Microbe 20, 392–405, doi:10.1016/j.chom.2016.08.005 (2016).

25 Schoggins, J. W. et al. A diverse range of gene products are effectors of the type I interferon antiviral response. Nature 472, 481–485, doi:10.1038/nature09907 (2011).

26 Feeley, E. M. et al. IFITM3 inhibits influenza A virus infection by preventing cytosolic entry. PLoS Pathog 7, e1002337, doi:10.1371/journal.ppat.1002337 (2011).

27 Staeheli, P., Grob, R., Meier, E., Sutcliffe, J. G. & Haller, O. Influenza virus-susceptible mice carry Mx genes with a large deletion or a nonsense mutation. Mol Cell Biol 8, 4518–4523, doi:10.1128/mcb.8.10.4518-4523.1988 (1988).

28 Verhelst, J., Parthoens, E., Schepens, B., Fiers, W. & Saelens, X. Interferon-inducible protein Mx1 inhibits influenza virus by interfering with functional viral ribonucleoprotein complex assembly. J Virol 86, 13445–13455, doi:10.1128/jvi.01682-12 (2012).

29 Wellington, D., Laurenson-Schafer, H., Abdel-Haq, A. & Dong, T. IFITM3: How genetics influence influenza infection demographically. Biomed J 42, 19–26, doi:10.1016/j.bj.2019.01.004 (2019).

30 Rhodes, D. A., Stammers, M., Malcherek, G., Beck, S. & Trowsdale, J. The cluster of BTN genes in the extended major histocompatibility complex. Genomics 71, 351–362, doi:10.1006/geno.2000.6406 (2001).

31 Kumar, S., Stecher, G., Suleski, M. & Hedges, S. B. TimeTree: A Resource for Timelines, Timetrees, and Divergence Times. Mol Biol Evol 34, 1812–1819, doi:10.1093/molbev/msx116 (2017).

32 Afrache, H., Pontarotti, P., Abi-Rached, L. & Olive, D. Evolutionary and polymorphism analyses reveal the central role of BTN3A2 in the concerted evolution of the BTN3 gene family. Immunogenetics 69, 379–390, doi:10.1007/s00251-017-0980-z (2017).

33 Pinto, R. M., Lycett, S., Gaunt, E. & Digard, P. Accessory Gene Products of Influenza A Virus. Cold Spring Harbor perspectives in medicine 11, doi:10.1101/cshperspect.a038380 (2021).

34 Naffakh, N., Tomoiu, A., Rameix-Welti, M. A. & van der Werf, S. Host restriction of avian influenza viruses at the level of the ribonucleoproteins. Annu Rev Microbiol 62, 403–424, doi:10.1146/annurev.micro.62.081307.162746 (2008).

35 Patrono, L. V. et al. Archival influenza virus genomes from Europe reveal genomic variability during the 1918 pandemic. Nat Commun 13, 2314, doi:10.1038/s41467-022-29614-9 (2022).

36 van Lieshout, L. P. et al. A Novel Triple-Mutant AAV6 Capsid Induces Rapid and Potent Transgene Expression in the Muscle and Respiratory Tract of Mice. Mol Ther Methods Clin Dev 9, 323–329, doi:10.1016/j.omtm.2018.04.005 (2018).

37 Portela, A. & Digard, P. The influenza virus nucleoprotein: a multifunctional RNA-binding protein pivotal to virus replication. J Gen Virol 83, 723–734, doi:10.1099/0022-1317-83-4-723 (2002).

38 Gabriel, G., Herwig, A. & Klenk, H. D. Interaction of polymerase subunit PB2 and NP with importin alpha1 is a determinant of host range of influenza A virus. PLoS Pathog 4, e11, doi:10.1371/journal.ppat.0040011 (2008).

39 Hu, Y., Sneyd, H., Dekant, R. & Wang, J. Influenza A Virus Nucleoprotein: A Highly Conserved Multi-Functional Viral Protein as a Hot Antiviral Drug Target. Curr Top Med Chem 17, 2271–2285, doi:10.2174/1568026617666170224122508 (2017).

40 Harly, C. et al. Key implication of CD277/butyrophilin-3 (BTN3A) in cellular stress sensing by a major human γδ T-cell subset. Blood 120, 2269–2279, doi:10.1182/blood-2012-05-430470 (2012).

41 Arnett, H. A. & Viney, J. L. Immune modulation by butyrophilins. Nature reviews. Immunology 14, 559–569, doi:10.1038/nri3715 (2014).

42 Gu, S., Borowska, M. T., Boughter, C. T. & Adams, E. J. Butyrophilin3A proteins and Vγ9Vδ2 T cell activation. Semin Cell Dev Biol 84, 65–74, doi:10.1016/j.semcdb.2018.02.007 (2018).

43 Galão, R. P. et al. TRIM25 and ZAP target the Ebola virus ribonucleoprotein complex to mediate interferon-induced restriction. PLoS Pathog 18, e1010530, doi:10.1371/journal.ppat.1010530 (2022).

44 Kuroda, M. et al. Identification of interferon-stimulated genes that attenuate Ebola virus infection. Nature Communications 11, 2953, doi:10.1038/s41467-020-16768-7 (2020).

45 Robinson, J. L. et al. Swine influenza (H3N2) infection in a child and possible community transmission, Canada. Emerg Infect Dis 13, 1865–1870, doi:10.3201/eid1312.070615 (2007).

46 Subbarao, E. K., London, W. & Murphy, B. R. A single amino acid in the PB2 gene of influenza A virus is a determinant of host range. J Virol 67, 1761–1764, doi:10.1128/jvi.67.4.1761-1764.1993 (1993).

47 Mehle, A. & Doudna, J. A. Adaptive strategies of the influenza virus polymerase for replication in humans. Proc Natl Acad Sci U S A 106, 21312–21316, doi:10.1073/pnas.0911915106 (2009).

48 Philippon, D. A. M., Wu, P., Cowling, B. J. & Lau, E. H. Y. Avian Influenza Human Infections at the Human-Animal Interface. J Infect Dis 222, 528–537, doi:10.1093/infdis/jiaa105 (2020).

49 Wang, D., Zhu, W., Yang, L. & Shu, Y. The Epidemiology, Virology, and Pathogenicity of Human Infections with Avian Influenza Viruses. Cold Spring Harbor perspectives in medicine 11, doi:10.1101/cshperspect.a038620 (2021).

50 Adlhoch, C., et al. Avian influenza overview December 2021 - March 2022. Efsa j 20, e07289, doi:10.2903/j.efsa.2022.7289 (2022).

51 Oliver, I. et al. A case of avian influenza A(H5N1) in England, January 2022. Euro Surveill 27, doi:10.2807/1560-7917.Es.2022.27.5.2200061 (2022).

52 Ramirez, R. D. et al. Immortalization of human bronchial epithelial cells in the absence of viral oncoproteins. Cancer Res 64, 9027–9034, doi:10.1158/0008-5472.Can-04-3703 (2004).

53 Wit, E. d., et al. Efficient generation and growth of influenza virus A/PR/8/34 from eight cDNA fragments. Virus Research 103, 155–161, doi:https://doi.org/10.1016/j.virusres.2004.02.028 (2004).

54 Rihn, S. J. et al. TRIM69 Inhibits Vesicular Stomatitis Indiana Virus. J Virol 93, doi:10.1128/JVI.00951-19 (2019).

55 Rihn, S. J. et al. A plasmid DNA-launched SARS-CoV-2 reverse genetics system and coronavirus toolkit for COVID-19 research. PLoS Biol 19, e3001091, doi:10.1371/journal.pbio.3001091 (2021).

56 Kawakami, E. et al. Strand-specific real-time RT-PCR for distinguishing influenza vRNA, cRNA, and mRNA. J Virol Methods 173, 1–6, doi:10.1016/j.jviromet.2010.12.014 (2011).

57 Bakshi, S., Taylor, J., Strickson, S., McCartney, T. & Cohen, P. Identification of TBK1 complexes required for the phosphorylation of IRF3 and the production of interferon β. Biochem J 474, 1163–1174, doi:10.1042/bcj20160992 (2017).

58 Balazs, A. B. et al. Antibody-based protection against HIV infection by vectored immunoprophylaxis. Nature 481, 81–84, doi:10.1038/nature10660 (2011).

59 Zufferey, R., Donello, J. E., Trono, D. & Hope, T. J. Woodchuck hepatitis virus posttranscriptional regulatory element enhances expression of transgenes delivered by retroviral vectors. J Virol 73, 2886–2892, doi:10.1128/jvi.73.4.2886-2892.1999 (1999).

60 Rghei, A. D. et al. Production of Adeno-Associated Virus Vectors in Cell Stacks for Preclinical Studies in Large Animal Models. J Vis Exp, doi:10.3791/62727 (2021).

61 MacLeod, M. K. et al. Vaccine adjuvants aluminum and monophosphoryl lipid A provide distinct signals to generate protective cytotoxic memory CD8 T cells. Proc Natl Acad Sci U S A 108, 7914–7919, doi:10.1073/pnas.1104588108 (2011).

62 Camacho, C., et al. BLAST+: architecture and applications. BMC Bioinformatics 10, 421, doi:10.1186/1471-2105-10-421 (2009).

63 Cunningham, F., et al. Ensembl 2022. Nucleic Acids Res 50, D988–d995, doi:10.1093/nar/gkab1049 (2022).

64 Mistry, J., Finn, R. D., Eddy, S. R., Bateman, A. & Punta, M. Challenges in homology search: HMMER3 and convergent evolution of coiled-coil regions. Nucleic Acids Res 41, e121, doi:10.1093/nar/gkt263 (2013).

65 Katoh, K. & Standley, D. M. MAFFT multiple sequence alignment software version 7: improvements in performance and usability. Mol Biol Evol 30, 772–780, doi:10.1093/molbev/mst010 (2013).

66 Suyama, M., Torrents, D. & Bork, P. PAL2NAL: robust conversion of protein sequence alignments into the corresponding codon alignments. Nucleic Acids Res 34, W609–612, doi:10.1093/nar/gkl315 (2006).

67 Nguyen, L. T., Schmidt, H. A., von Haeseler, A. & Minh, B. Q. IQ-TREE: a fast and effective stochastic algorithm for estimating maximum-likelihood phylogenies. Mol Biol Evol 32, 268–274, doi:10.1093/molbev/msu300 (2015).

68 Steinegger, M. & Söding, J. MMseqs2 enables sensitive protein sequence searching for the analysis of massive data sets. Nat Biotechnol 35, 1026–1028, doi:10.1038/nbt.3988 (2017).

69 Sagulenko, P., Puller, V. & Neher, R. A. TreeTime: Maximum-likelihood phylodynamic analysis. Virus Evol 4, vex042, doi:10.1093/ve/vex042 (2018).

70 Hoang, D. T., Chernomor, O., von Haeseler, A., Minh, B. Q. & Vinh, L. S. UFBoot2: Improving the Ultrafast Bootstrap Approximation. Mol Biol Evol 35, 518–522, doi:10.1093/molbev/msx281 (2018).

71 Yu, G. Using ggtree to Visualize Data on Tree-Like Structures. Curr Protoc Bioinformatics 69, e96, doi:10.1002/cpbi.96 (2020).

72 Huerta-Cepas, J., Serra, F. & Bork, P. ETE 3: Reconstruction, Analysis, and Visualization of Phylogenomic Data. Mol Biol Evol 33, 1635–1638, doi:10.1093/molbev/msw046 (2016).

