## Supplementary Figures for "Zoonotic avian influenza viruses evade human BTN3A3 restriction"

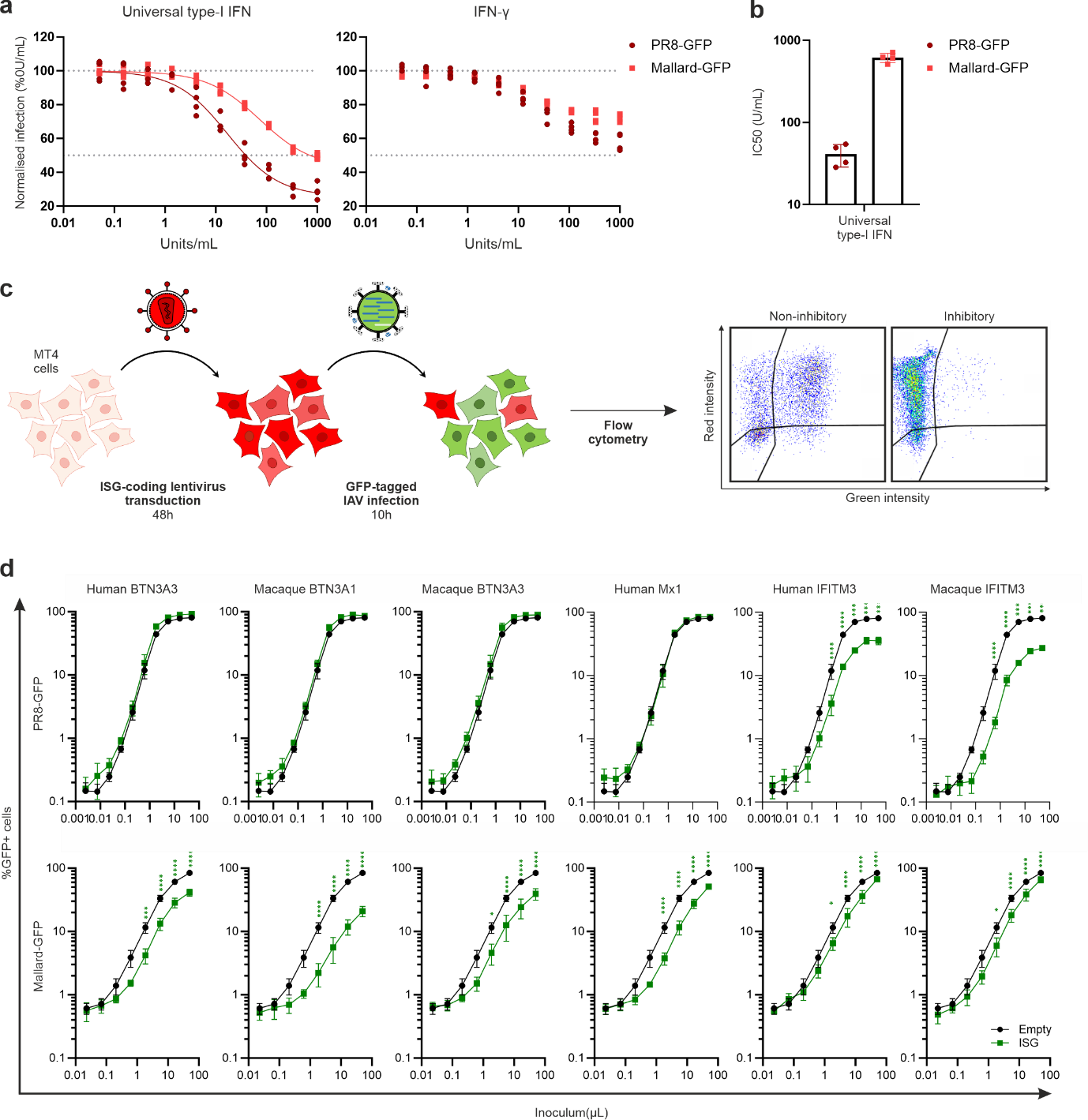

**Extended Data Figure 1: Arrayed ISG expression screening and validation data. a**, Dose-response curves assessing the ability of type-I and type-III IFN to protect A549 cells from PR8-GFP and Mallard-GFP. **b,** Estimated IC_50_ from data in a. **c**, Diagrammatic representation of the ISG screening method used in Figure 1a-b. **d**, Validation data of the larger screens shown in Fig. 1b. MT4 cells were transiently transduced with SCPRSY lentiviral vectors encoding the indicated human or macaque ISGs. 48h post-transduction, cells were infected with serially diluted GFP-tagged PR8 or Mallard for 10h. Cells were fixed and analysed by flow cytometry. Data are mean +/- SEM of 3 independent experiments. Statistical significance between groups was measured by a 1-way ANOVA for each virus dose. *p≤0.05, **p≤0.01, ***p≤0.001, ****p≤0.0001.

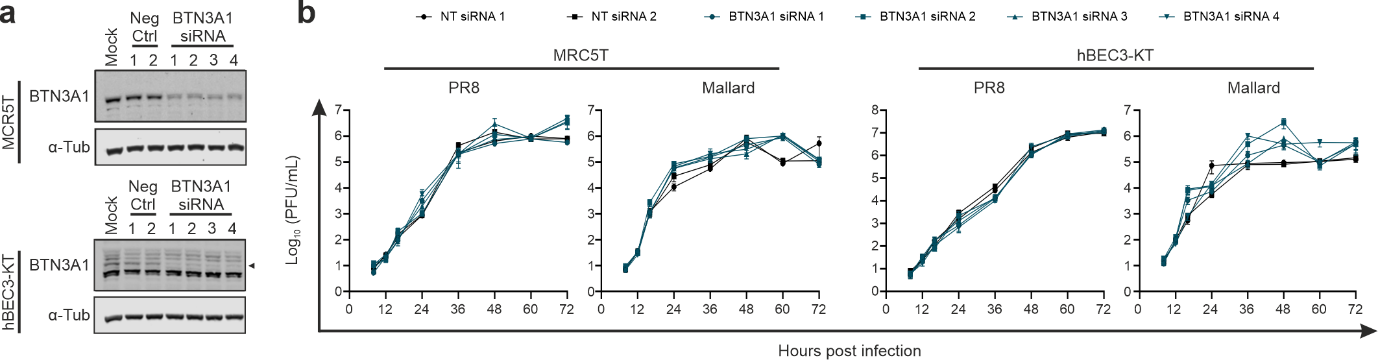

**Extended Data Fig. 2: Limited antiviral effects of BTN3A1 in MRC5T and hBEC3-KT cells. a,** Western blotting showing the efficacy of siRNA knock-down of BTN3A1 in MRC5T and hBEC3-KT. Cells were transfected with scrambled (Neg ctrl) or BTN3A1-targetting siRNAs, and protein levels in the resulting cell lysates were assessed by western blotting. α-Tubulin was used as loading control. Arrows indicate the band corresponding to BTN3A1. **b**, Graphs showing the replication kinetics of PR8 and Mallard in siRNA-treated MRC5T and hBEC3-KT cells. Cells were infected with a MOI of 0.001 and supernatants were collected at the mentioned times post infection. Infectious viral titres were determined by plaque assay. Antiviral activity by BTN3A1 is less clear compared to the one displayed by BTN3A3 and showed in Fig. 1d-e. Data are mean +/- SEM of 3 independent experiments. Statistical significance between groups was measured by a 2-way ANOVA. Comparisons were made between area under the curve of the different BTN3A1 siRNA treatment conditions and the average of the two negative controls. No statistically significant differences were found.

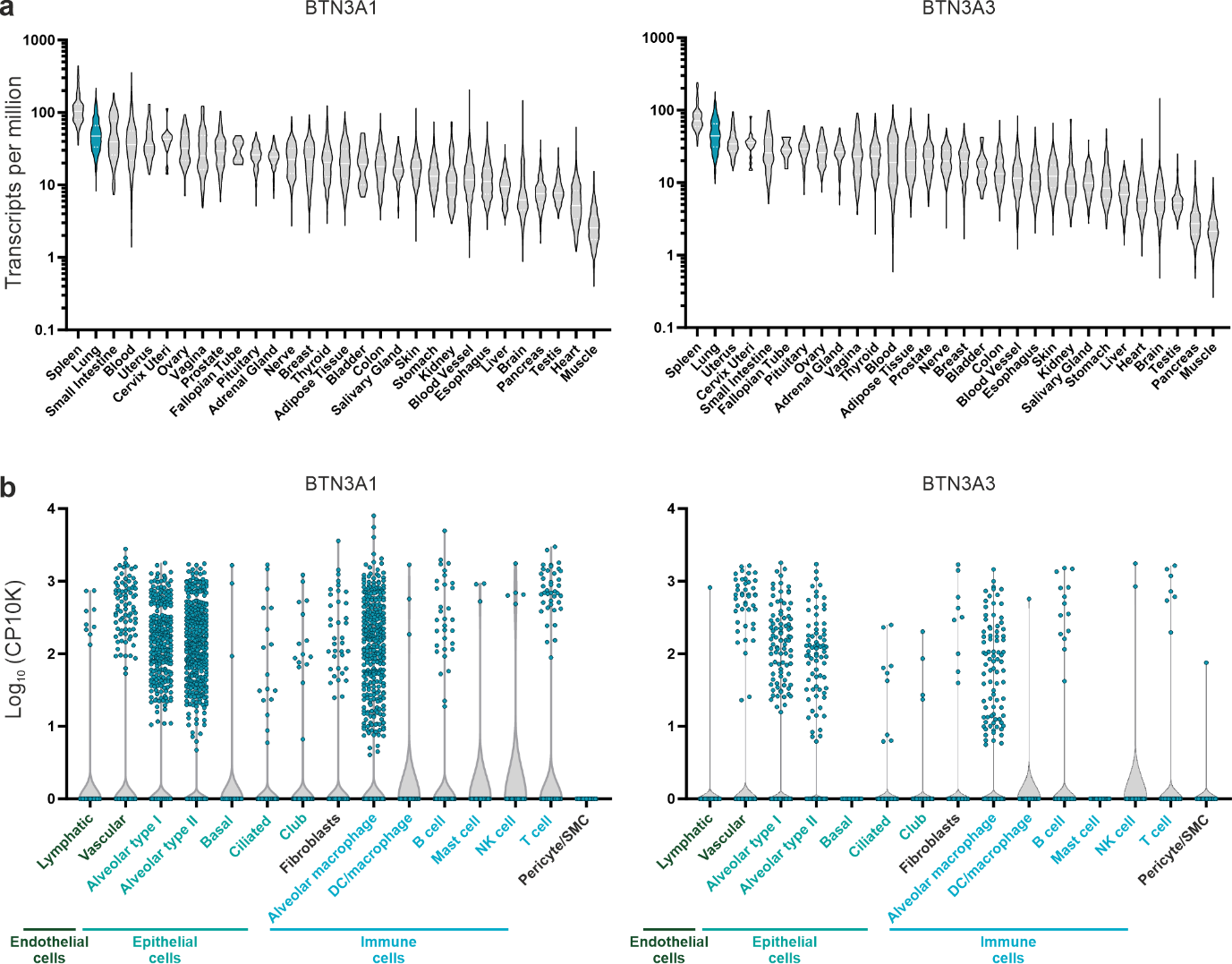

**Extended Data Fig. 3: BTN3A1 and BTN3A3 gene expression across a panel of tissues. a**, Organ-dependent bulk tissue gene expression. **b**, Lung single cell tissue expression. Data was obtained from the GTEx Portal ([www.gtexportal.org](http://www.gtexportal.org)).

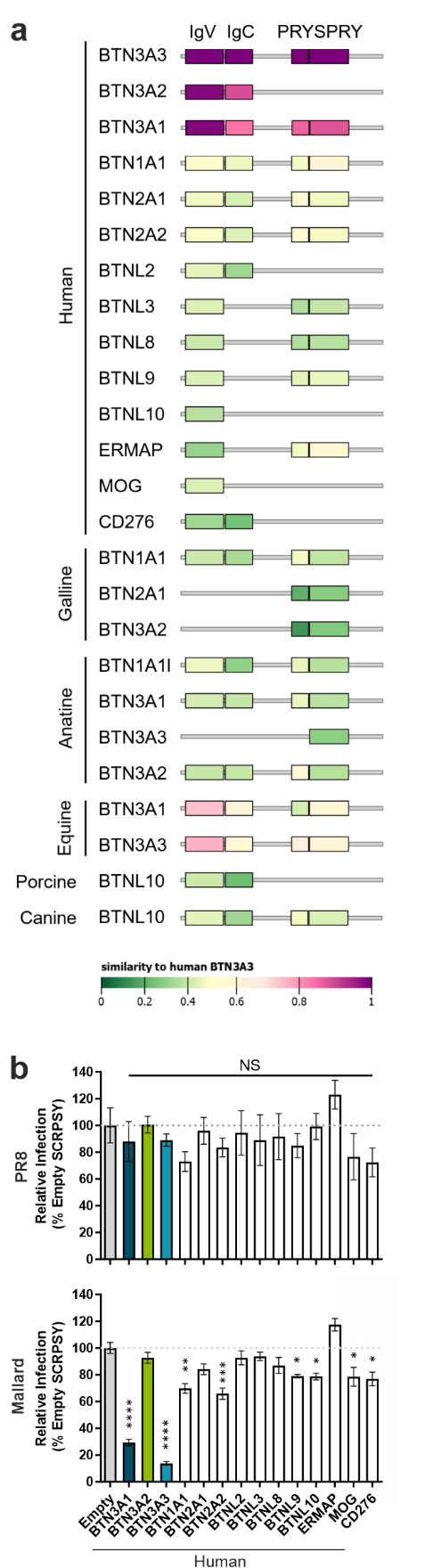
**Extended Data Figure 4: Domain organisation and antiviral activity of BTN orthologous and paralogues.** **a**, Schematic representation of domain organisation and sequence similarity of the indicated proteins. **b**, A549 cells were transiently transduced with SCRPSY lentiviruses expressing the indicated human BTN proteins and challenged with PR8- or Mallard-GFP. 8 h post infection, percentage of GFP-positive cells was measured by flow cytometry. Data are mean +/- SEM of 2 independent experiments. Statistical significance between groups was measured by a 1-way ANOVA. Comparisons were made between each BTN-expressing cells and empty control. NS-Non-significant, *p≤0.05, **p≤0.01, ***p≤0.001, ****p≤0.0001.

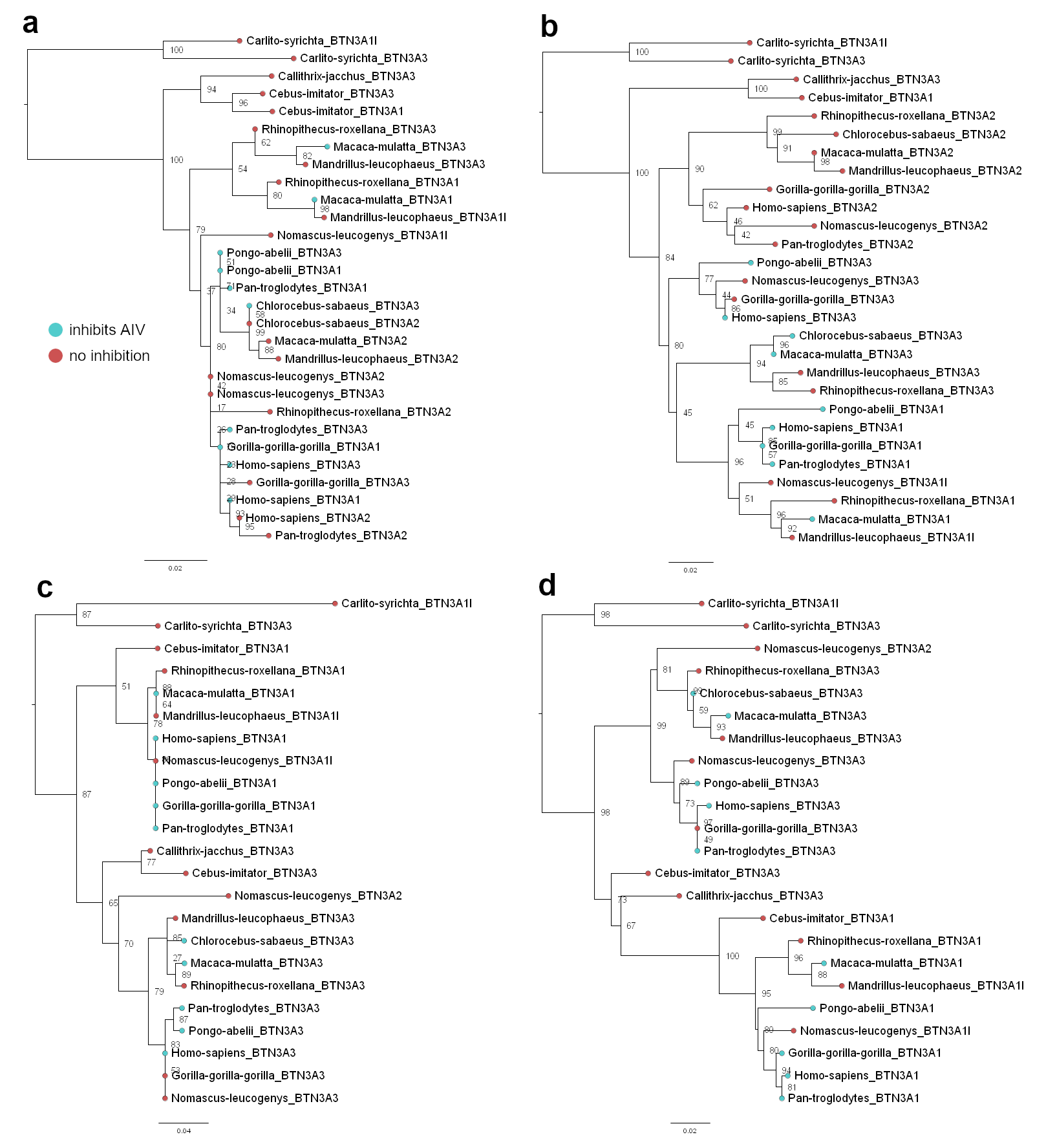

**Extended Data Figure 5: Maximum likelihood *Haplorrhini* BTN3 gene coding sequence phylogenies of separate domains**: IgV (a), IgC (b), PRY (c), SPRY (d) under a K2P+G4 substitution model. Trees are rooted at the *C. syrichta* branch and node confidence values (10,000 bootstrap replicates) are annotated on each node. Tip shapes are coloured by whether each gene exhibits anti-AIV activity (consistent with Fig. 3a). Phylogenies were visualised using FigTree.

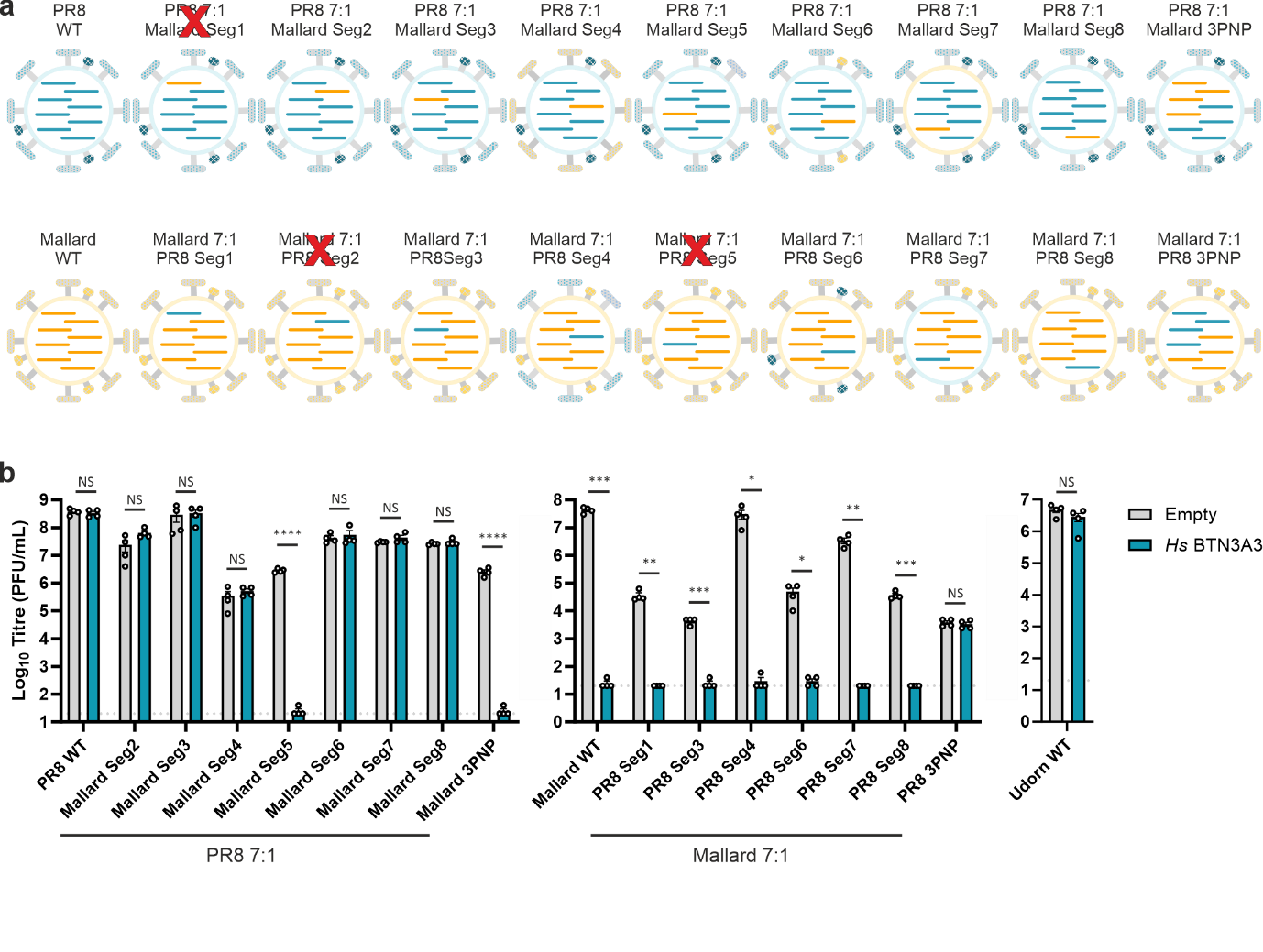

**Extended data Figure 6: BTN3A3 sensitivity of PR8 - Mallard reassortants. a**, Schematic representation of reassortant viruses between PR8 (blue) and Mallard (yellow) viruses. Crossed titles represent viruses that have failed to rescue (PR8 7:1 Mallard seg 1, Mallard 7:1 PR8 seg 2, Mallard 7:1 PR8 seg 5). **b**, PR8:Mallard reassortants represented in A and Udorn WT were used to perform plaque assays in MDCK cells expressing BTN3A3. Data are mean +/- SEM of 2 technical replicates from 2 independent experiments. Statistical differences between Empty and *Hs*BTN3A3 were calculated using multiple t-tests and corrected for multiple comparisons using the Holm-Šídák method. NS-Non-significant, *p≤0.05, **p≤0.01, ***p≤0.001, ****p≤0.0001.

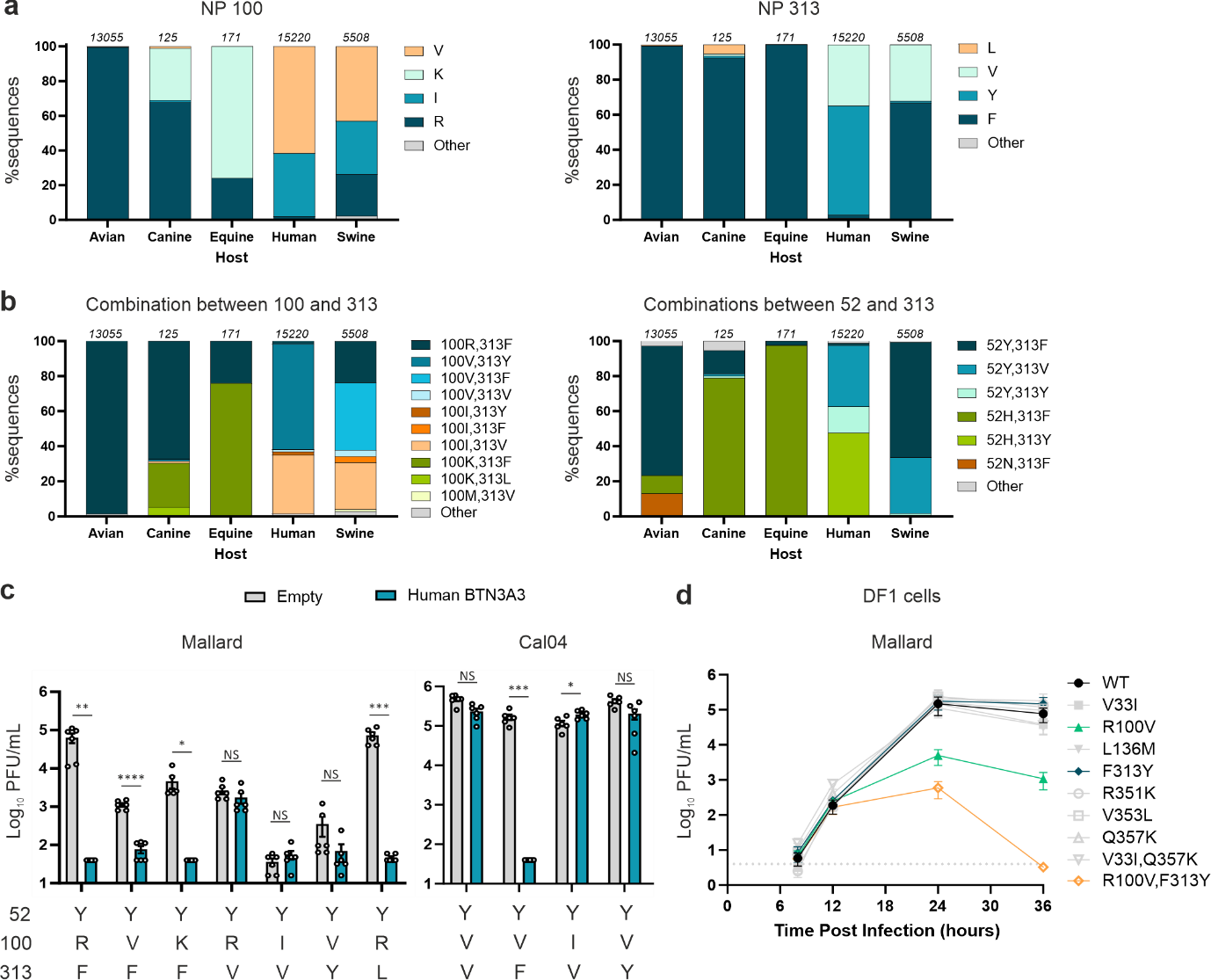

**Extended Data Figure 7: Variation of residues 52, 100 and 313, BTN3A3-sensitivity and viral fitness. a**, Identity of amino acid residues in positions 100 (left) and 313 (right) of all NP proteins available in our dataset. **b**, Identity of combinations between amino acids 100 and 313 (left) and 52 and 313 (right). **c**, Infectious virus titres obtained in A549-Empty and A549-BTN3A3 cells infected with either Mallard or Cal04 residues 100 and 313 mutants. Cells were infected with an MOI of 0.001 for 48 h. Infectious virus titres were acquired by plaque assay. Data are mean +/- SEM of 2 technical replicates from 3 independent experiments. Statistical differences between cells expressing BTN3A3 and control cells were calculated using multiple t-tests and corrected for multiple comparisons using the Holm-Šídák method. NS-Non-significant, *p≤0.05, **p≤0.01, ***p≤0.001, ****p≤0.0001. **d**, Viral replication assays in avian cells were carried out in chicken fibroblasts (DF1 cells). Cells were infected an MOI of 0.001 for 48 hours. Infectious virus titres were determined by plaque assay.

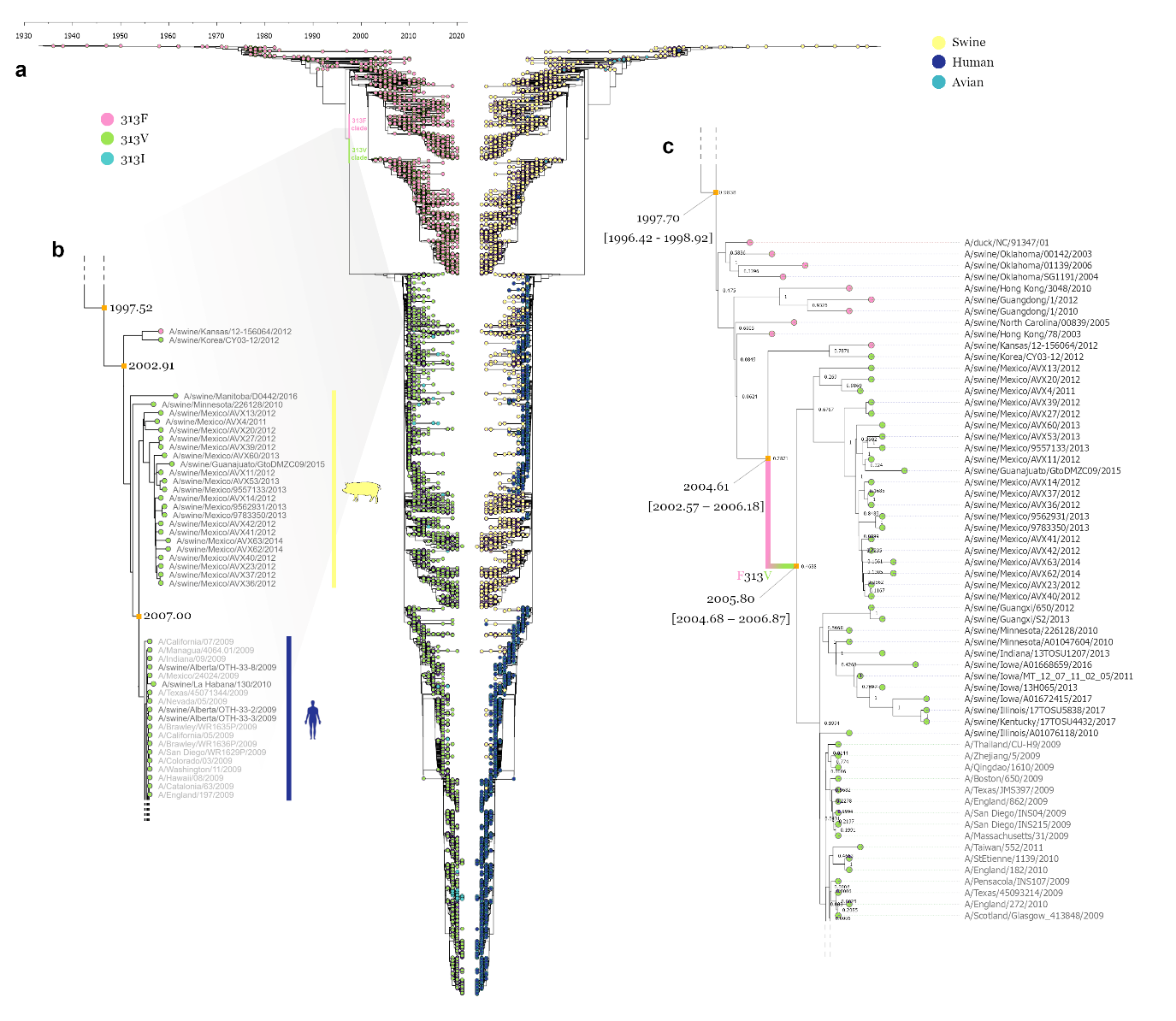

**Extended Data Figure 8. Molecular dating of the F313V NP substitution on the classical swine H1N1 lineage. a**, Tip-dated maximum likelihood phylogeny of all classical H1N1 lineage NP sequences annotated by position 313 residue (left) and isolation host (mirrored tree, right). **b**, Zoomed in snippet of the part of the ML phylogeny shown in A where the F313V change has occurred. Tip shapes are coloured by 313 residue, estimated dates for key nodes are annotated, and strain names are shown on the right of the tips. **c**, Zoomed in snippet of the part of the BEAST maximum clade credibility phylogeny where the F313V change has occurred. Tip shapes are coloured by 313 residue, median node age and 95% highest posterior density confidence intervals are annotated for key nodes, posterior probability values are shown for each node, and strain names are shown on the right of the tips. The branch where F313V is believed to have taken place on is annotated in colour (pink and green). Phylogenies were visualised using FigTree.

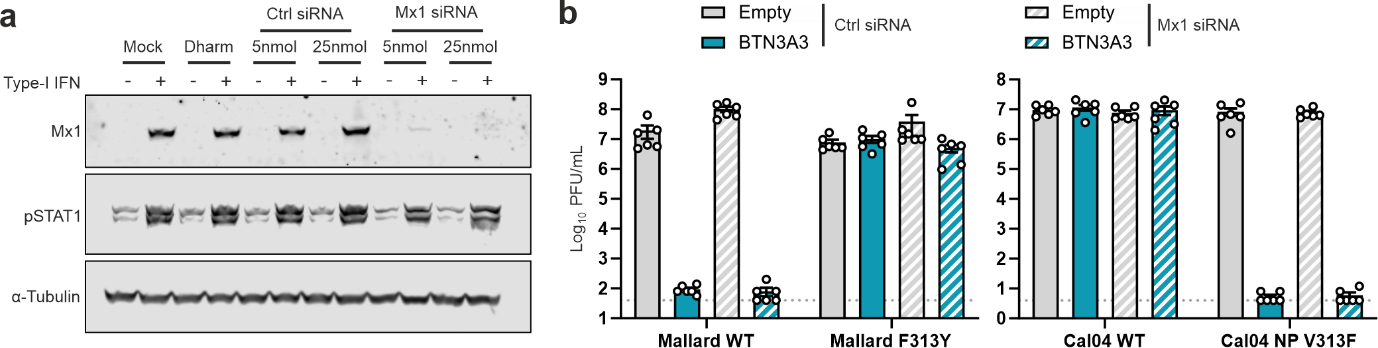

**Extended Data Fig. 9: BTN3A3 activity is independent from Mx1. a**, Western blotting of cell lysates obtained from A549 cells transfected with either control siRNAs or siRNA pools targeting Mx1 siRNA. Cells were transfected for 48h followed by treatment for 16h with type-I IFN. pSTAT1 and α-Tubulin were used as IFN-treatment and loading controls, respectively. **b**, Infectious virus assays. Upon siRNA treatment, A549 Empty and BTN3A3 cells were infected with the indicated viruses at an MOI of 0.001. Supernatants were harvested at 48 hpi and infectious viral titres were measured by plaque assay. Knock down of Mx-1 does not affect BTN3A3 restriction. Data are mean +/- SEM of 2 technical replicates from 3 independent experiments.

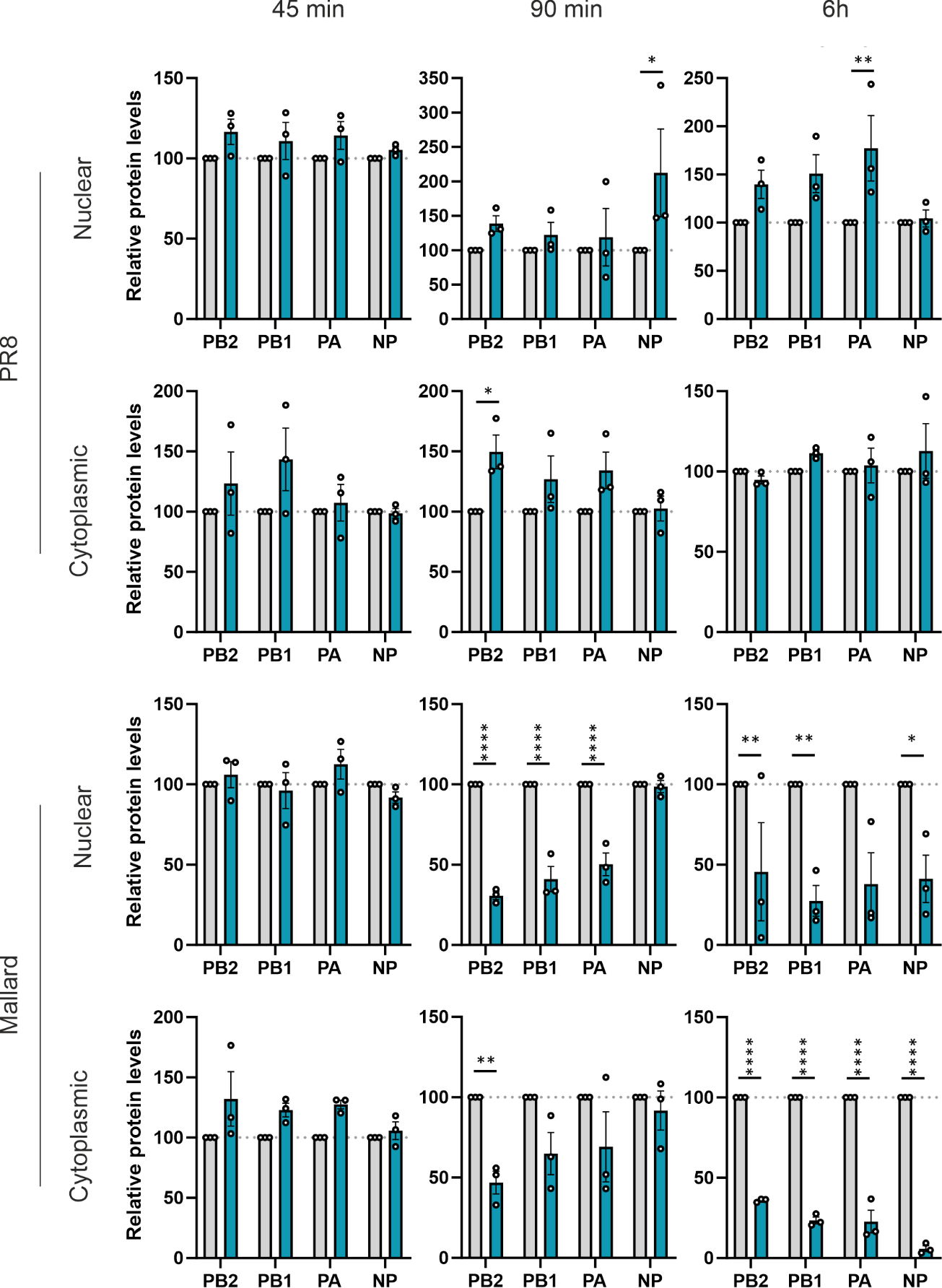

**Extended Data Figure 10. Quantification of cytoplasmic and nuclear levels of vRNP complex proteins at early stages post infection in the presence or absence of BTN3A3.** Quantification of three independent western blots, one set of which is shown in Fig. 5a. A549-Empty and BTN3A3 overexpressing cells were synchronously infected with PR8 or Mallard at MOI 3. Nuclear/cytoplasm fractionation was performed at 45, 90 mins and 6h post infection. Quantification of vRNP-complex proteins was performed by fluorescence. Cytoplasmic and nuclear viral proteins were normalised to GAPDH and H3, respectively. All values were further normalised to what obtained in A549-Empty cells. Data are mean +/- SEM of 3 independent experiments. Statistical significance between groups was measured by a 2-way ANOVA. Comparisons were made between A549-Empty and A549-BTN3A3. NS- non-significant, * p ≤ 0.05, ** p ≤ 0.01, *** p ≤ 0.001, **** p ≤ 0.0001.

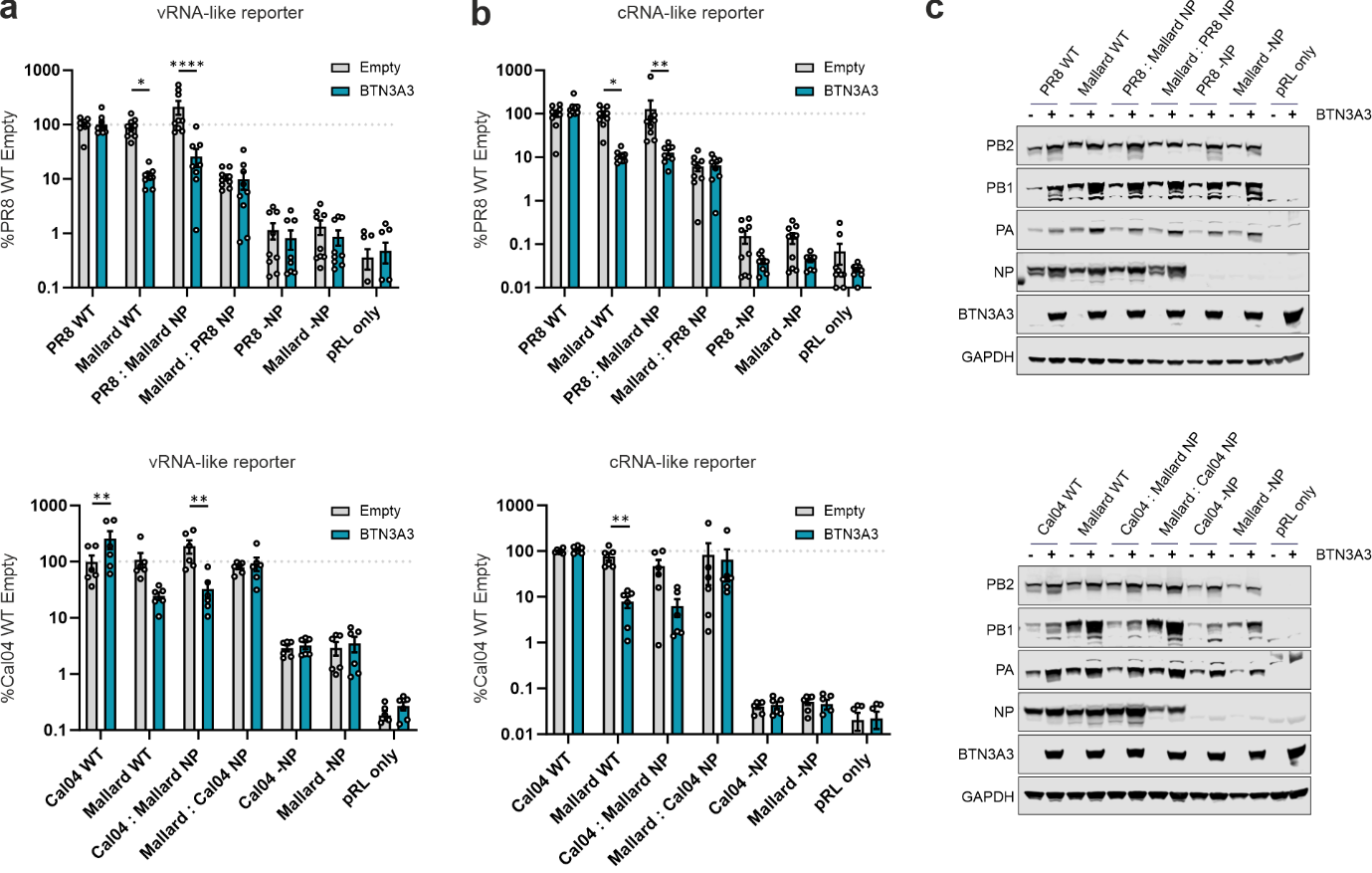

**Extended Data Fig. 11: Minireplicon assays with avian/mammalian NP reassortant RNP complexes**. **a-b**, A549-Empty and A549-BTN3A3 cells were transfected with pDUAL plasmids encoding for PB2, PB1, PA and NP of the indicated viruses alongside firefly luciferase-coding vRNA- (a) or cRNA-like (b) reporter plasmids. Forty-eight hours post transduction, cells were lysed and firefly luciferase was measured. Values were normalised to PR8 or Mallard WT replicons without NP (PR8 -NP; Mallard -NP) transfected in A549-Empty. Data are mean +/- SEM of 3 technical replicates from 3 independent experiments. Statistical differences between Empty and *Hs*BTN3A3 were calculated using multiple t-tests and corrected for multiple comparisons using the Holm-Šídák method. NS-Non-significant, *p≤0.05, **p≤0.01, ***p≤0.001, ****p≤0.0001. **c**. Expression levels of PB2, PB1, PA and NP from a and b were confirmed by western blot (GAPDH was used as loading control).

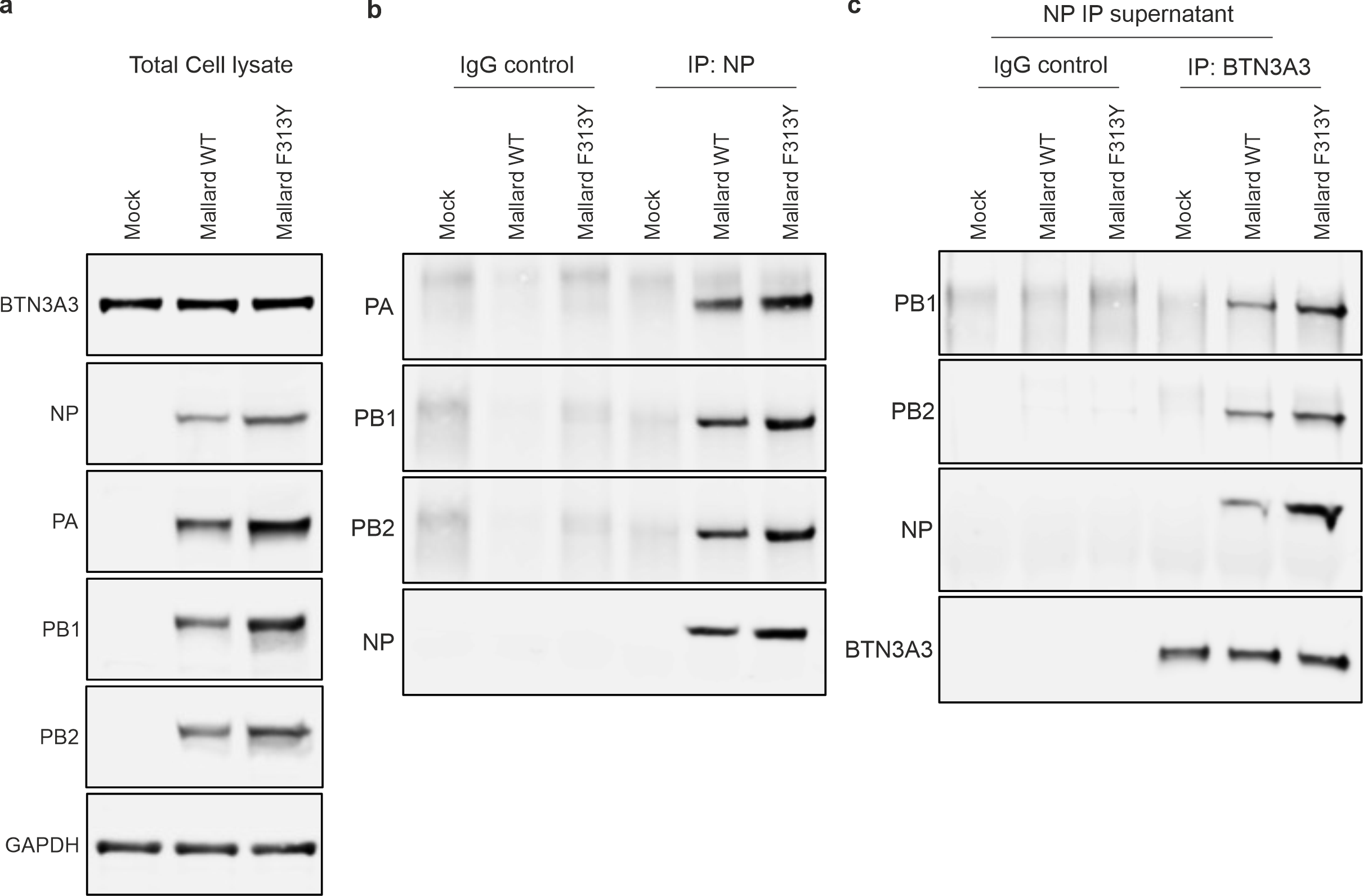

**Extended Data Figure 12: Interactions between NP and BTN3A3.** **a**, hBEC3-KT cells were infected with IAV Mallard or the the equivalent BTN3A3-resistant mutant (Mallard NP F313Y) virus for 6 hours and western blotting of NP, PA, PB1, PB2 and GAPDH (loading control) from total cell lysates is shown. **b**, Immunoprecipitation of NP pulls down not only NP but also the remaining protein components of the RNP complex. **c**, Supernatant of the NP immunoprecipitates was used to perform an additional immunoprecipitation using an anti-BTN3A3 antibody and PB2, PB1 and NP were amongst the co-immunoprecipitates.

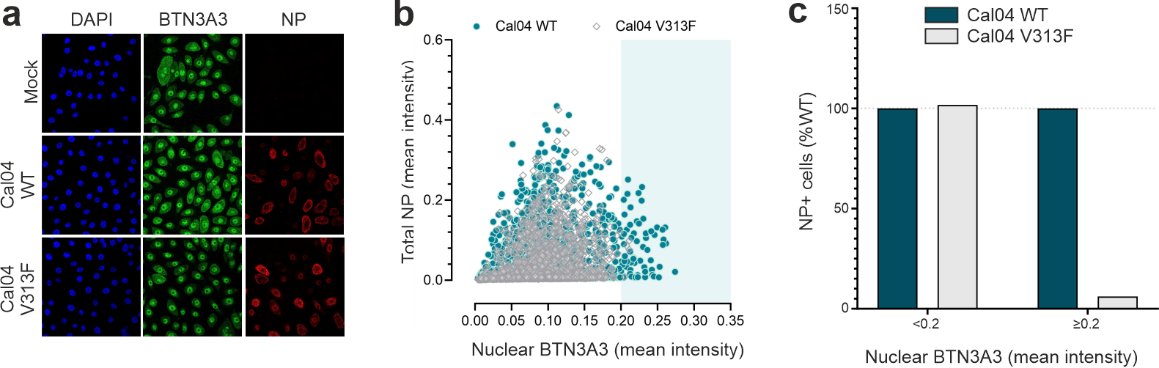

**Extended Data Figure 13. Quantification of the correlation between NP signal and nuclear BTN3A3. a,** Representative image of confocal microscopy of hBEC3-KT cells infected with Cal04WT or Cal04 NP V313F at MOI 3. Six hours post infection, cells were immunostained with NP (red) and BTN3A3 (green). DAPI staining (blue) was used as a nuclear marker. Image acquisition was performed with a Zeiss 880 confocal microscope using a 63x objective. **b**, Images from >3500 cells from four independent experiments performed as in (a) were used to quantify total NP and nuclear BTN3A3 for Cal04 WT and Cal04 NP V313F **c**, Values from b were stratified based on nuclear BTN3A3 intensity (<0.2 or ≥0.2). Data represents relative abundance of total infected cells present in each of the two nuclear BTN3A3 intensities ranges, taking values obtained with Cal04 WT as 100%.

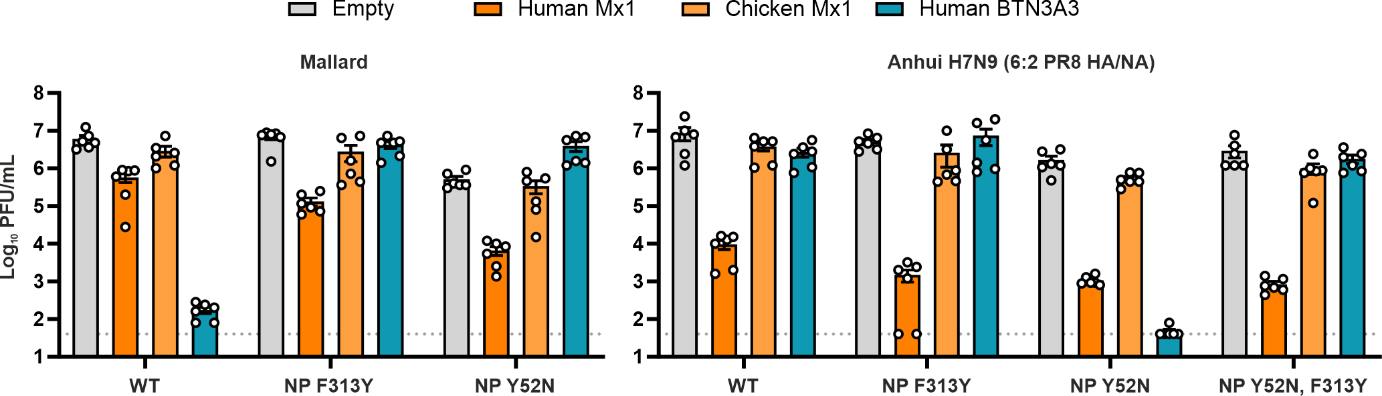

**Extended Data Fig. 14: Differential requirements of NP residues 52 and 313 for BTN3A3 and Mx1 sensitivity and evasion.** A549 cells overexpressing human or galline Mx1, or human BTN3A3 were infected with the mentioned viruses at a MOI of 0.001. Supernatants were harvested at 48 hpi and infectious viral titres were measured by plaque assay. Data are mean +/- SEM of 2 technical replicates from 3 independent experiments.

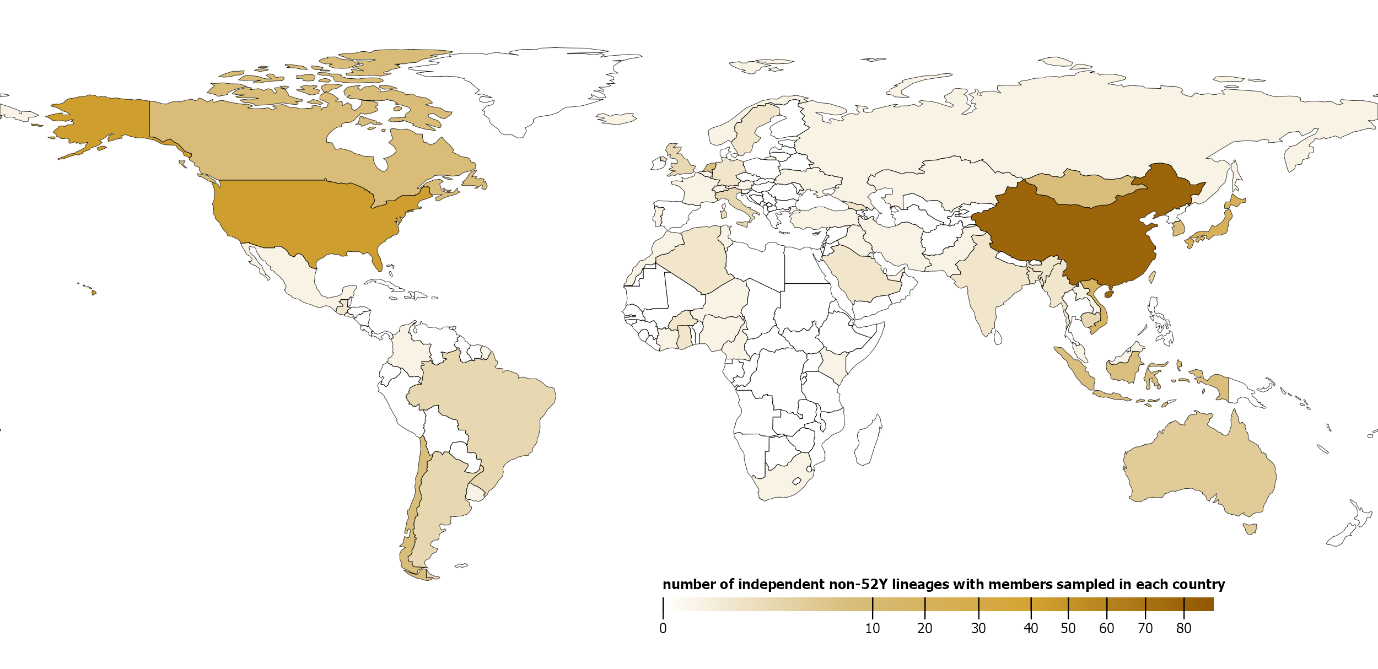

**Extended Data Fig. 15: Geographical distribution of the independent lineages of the avian clade with a BTN3A3-resistant genotype.** 151 independent lineages (each one consisting of at least two members) were identified with no 52Y residues in the avian clade of the IAV NP phylogeny presented in Fig. 4c. Map shading corresponds to the number of these lineages that each country has sampled at least one isolate from.

**Supplementary Tables Legends**

Table S1. NCBI accessions of all BTN3 homologues presented in this study.

Table S2. Classical H1N1 swine clade NP sequence NCBI accessions with sub-clade and BEAST dataset annotations.

Table S3. IAV isolates sampled in avian hosts in Europe between the 1st of January 2021 and the 11th of June 2022 available on the GISAID database. Residues at positions 52 and 313 are annotated for each accession.

Table S4. Proportions of avian IAV isolates with H, N and Y 52 residues by country, sampled in Europe between the 1st of January 2021 and the 11th of June 2022.

Table S5. IAV H5 isolates sampled in mammalian hosts (including humas) between the 1^st^ of January 2021 and the 11^th^ of June 2022 available on the GISAID database. Residues at positions 52 and 313 are annotated for each accession.

Table S6. GISAID EpiFlu acknowledgments for the isolates used in this study.
