## Supplementary Text 1 for "Zoonotic avian influenza viruses evade human BTN3A3 restriction"

**Text S1: Molecular dating of the F313V NP change**

Based on the tree topology of the MMseqs2 filtered dataset including sequences from all IAV NP clades, representative sequences from the classical swine H1N1 clade as well as the unfiltered sequences from each representative’s corresponding cluster were retrieved (9,426 sequences, see Methods). The codon alignment of all swine clade NP sequences was used to infer a more detailed maximum likelihood phylogenetic reconstruction of this particular clade (iqtree under a GTR+I+F+G4 model with 10,000 Ultrafast bootstrap replicates)^1,2^. The phylogeny was time-calibrated using TreeTime^3^ as described in the ‘IAV phylogenetic analysis’ section (Extended Data Fig. 8a).

The phylogeny reveals a clear split between the 313F and 313V clades with a relatively long branch consistent with the F313V change. The divergence date between the two major 313F/V clades is estimated to be 1997.52. Upon closer inspection, two isolates: A/swine/Korea/CY03-12/2012(H3N1) (AGE83887) and A/swine/Kansas/12-156064/2012(H1N2) (AHN19642) sampled in South Korea and the US respectively^4,5^, are the most basal within the 313V clade. Interestingly, the NP of the latter isolate has a 313F residue. Assuming that this topology is true, the most parsimonious explanation would be that the major F313V change took place after the split between this two-isolate clade and the 313V clade, while an independent F313V change occurred in the A/swine/Korea/CY03-12/2012 lineage. The alternative explanation would require a reversion back to 313F unique to the A/swine/Kansas/12-156064/2012 lineage. This means that we can narrow down the window of time when the F313V change occurred to start from an estimated date of 2002.91 (Extended Data Fig. 8b). Within the 313V clade a distinct subclade consisting of isolates sampled in swine hosts primarily from Mexico between 2011 and 2015 sits as outgroup to the H1N1 pdm09 NP sequences. All isolates in this subclade have a 313V residue, supporting a F313V change responsible for human butyrophilin antiviral evasion taking place in the swine host reservoir, prior to human emergence (Extended Data Fig. 8b). Furthermore, the 313 residue change should have happened prior to the date of the split between the swine subclade and the pdm09 lineage, estimated to be 2007.00.

Inferring phylogenies for large datasets such as this can sometimes lead to erroneous topologies, especially if sampling biases exist between different clades in the tree (for example there are many more sequences from the H1N1 pdm09 lineage compared to any other clade in the tree). To validate the inferred topology we used an independent Bayesian phylogenetic approach on a subset of the full dataset. From the original dataset (9,426 sequences), the sequences outwith the distinct 313F and 313V subclades (Extended Data Fig. 8a) and sequences topologically close to the F313V change branch - based on the maximum likelihood phylogeny (for example the swine 313V subclade) - were kept in the subset (263 sequences, see Table S2). To remove sampling biases from the two remaining subclades we used a subsampling approach where up to 200 sequences were retrieved from each subclade but a maximum of 200 divided by the number of sampling years were kept for each sampling year. For example, if a subclade had sequences from sampling years 2008, 2009, 2010 and 2011 with sequence counts 35, 1000, 800 and 1200 respectively, then all 35 sequences from 2008 would be kept and only 50 (200/4) from each of the other three years. This approach yielded a total of 380 subsampled sequences from the two 313F/V subclades and combined with the 263 non-subsampled isolates produced a total of 643 NP sequences to be analysed using Bayesian phylogenetics.

These sequences were retrieved from the full dataset codon NP alignment and used to infer a BEAST (v1.10.4) phylogeny under a HKY model, accounting for site heterogeneity with a 4 category Γ distribution^6^. Codon positions were evaluated separately by the model and sampling years were used for tip-dating. Two independent MCMC chains, 150,000,000 states long each, were performed, sampling every 150,000 states. LogCombiner was used to combine the two independent chains after removing 20% burn-in states from each chain, ensuring chain convergence and an effective sample size >200 for the joint parameters.

Despite low support clustering of a few more 313F sequences within the 313V clade, the aforementioned South Korea and US isolate clade is consistently found within the 313V clade (node posterior = 0.79, Extended Data Fig. 8c). The topology of the swine 313V clade from Mexico is also congruent with the maximum likelihood phylogeny. This independent validation of the topology inference strongly supports the above assessment of the F313V change taking place in swine hosts prior to human emergence. The BEAST phylogeny can also provide us with more accurate node dating estimates, placing the earliest date for the F313V change in 2004.61 (95% HPD: 2002.57 – 2006.18) and latest date in 2005.80 (95% HPD: 2004.68 – 2006.87). These estimates are largely consistent with the time-calibrated maximum likelihood phylogeny and support that F313V very likely occurred between mid-2002 and the end of 2006.

**Text S1 References**

1.	Nguyen, L. T., Schmidt, H. A., Von Haeseler, A. & Minh, B. Q. IQ-TREE: A fast and effective stochastic algorithm for estimating maximum-likelihood phylogenies. *Mol. Biol. Evol.* **32**, 268–274 (2015).

2.	Hoang, D. T., Chernomor, O., Von Haeseler, A., Minh, B. Q. & Vinh, L. S. UFBoot2: Improving the ultrafast bootstrap approximation. *Mol. Biol. Evol.* **35**, 518–522 (2018).

3.	Sagulenko, P., Puller, V. & Neher, R. A. TreeTime: Maximum-likelihood phylodynamic analysis. *Virus Evol.* **4**, (2018).

4.	Noriel, P. *et al.* Emergence of H3N2pM-like and novel reassortant H3N1 swine viruses possessing segments derived from the A (H1N1)pdm09 influenza virus, Korea. doi:10.1111/irv.12154.

5.	Duff, M. A. CHARACTERIZATION OF H1N2 VARIANT INFLUENZA VIRUSES IN PIGS. (Kansas State University, Manhattan, Kansas, 2014).

6.	Suchard, M. A. *et al.* Bayesian phylogenetic and phylodynamic data integration using BEAST 1.10. *Virus Evol.* **4**, (2018).
